# Microtubules originate asymmetrically at the somatic Golgi and are guided via Kinesin2 to maintain polarity within neurons

**DOI:** 10.1101/846832

**Authors:** Amrita Mukherjee, Paul Brooks, Fred Bernard, Antoine Guichet, Paul T. Conduit

## Abstract

Neurons contain polarised microtubule arrays essential for neuronal function. How microtubule nucleation and polarity are regulated within neurons remains unclear. We show that γ-tubulin localises asymmetrically to the somatic Golgi within *Drosophila* neurons. Microtubules originate from the Golgi with an initial growth preference towards the axon. Their growing plus ends also turn towards and into the axon, adding to the plus-end-out microtubule pool. Any plus ends that reach a dendrite, however, do not readily enter, maintaining minus-end-out polarity. Both turning towards the axon and exclusion from dendrites depend on Kinesin-2, a plus-end-associated motor that guides growing plus ends along adjacent microtubules. We propose that Kinesin-2 engages with a polarised microtubule network within the soma to guide growing microtubules towards the axon; while at dendrite entry sites engagement with microtubules of opposite polarity generates a backward stalling force that prevents entry into dendrites and thus maintains minus-end-out polarity within proximal dendrites.

## Introduction

Microtubules are polarised α/β-tubulin-based polymers essential for cell viability, providing pushing and pulling forces, structural support or tracks for the transport of intracellular cargo (Goodson and Jonasson, 2018). α-tubulin is located at the so-called minus end and β-tubulin is exposed at the so-called plus end, which is typically more dynamic. This inherent microtubule polarity is important for cell polarity, as different motor proteins (Kinesins and Dynein) move cargo along microtubules in a specific direction – Dynein towards the minus end and most Kinesins towards the plus end. In neurons, most plus ends point away from the soma in axons (plus-end-out microtubules), while the microtubules in dendrites are either of mixed polarity or are predominantly minus-end-out (Hill et al., 2012; Kapitein and Hoogenraad, 2015; Kelliher et al., 2019; Tas et al., 2017). This difference between axons and dendrites is important for the correct distribution of cargo throughout the neuron (Harterink et al., 2018; Kapitein and Hoogenraad, 2015; Tas et al., 2017).

Within cells *de novo* assembly of new microtubules, i.e. microtubule nucleation, is kinetically unfavourable and is templated and catalysed by multi-protein γ-tubulin ring complexes (γ-TuRCs) (Tovey and Conduit, 2018). Knockdown of γ-TuRCs within model systems affects dynamic microtubules in all neuronal compartments (Nguyen et al., 2014; Ori-McKenney et al., 2012; Sánchez-Huertas et al., 2016; Yamada and Hayashi, 2019; Yau et al., 2014) and mutations in γ-TuRC genes have been linked to human neurodevelopmental disorders (Bahi-Buisson et al., 2014; Mitani et al., 2019; Poirier et al., 2013). γ-TuRCs are typically inactive until they are recruited to specific sites within cells, such as microtubule organising centres (MTOCs), the cytosol around mitotic chromatin, or the sides of pre-existing microtubules via binding to Augmin/HAUS complexes (Farache et al., 2018; Lin et al., 2014; Meunier and Vernos, 2016; Sanchez and Feldman, 2016; Teixidó-Travesa et al., 2012). A range of MTOCs exist, including centrosomes, the Golgi apparatus and the nuclear envelope, and different cells use different MTOCs to help generate and organise their specific microtubule arrays (Sanchez and Feldman, 2016). γ-TuRC recruitment occurs via γ- TuRC “tethering proteins”, such as *Drosophila* Centrosomin (Cnn), that simultaneously bind to the γ-TuRC and a particular MTOC, and can also help activate the γ-TuRC (Tovey and Conduit, 2018).

Although γ-tubulin is important within neurons (Nguyen et al., 2014; Ori-McKenney et al., 2012; Sánchez-Huertas et al., 2016; Yamada and Hayashi, 2019; Yau et al., 2014), it remains unclear how microtubule nucleation is regulated. During early development of mammalian neurons, the centrosome within the soma nucleates microtubules (Stiess et al., 2010) that are severed and transported into neurites via motor-based microtubule sliding (Baas et al., 2005). Microtubule sliding is also important for axon outgrowth in *Drosophila* cultured neurons (Castillo et al., 2014; Lu et al., 2013), and for establishing microtubule polarity (Castillo et al., 2015; Klinman et al., 2017; Rao et al., 2017; Yan et al., 2013; Zheng et al., 2008). Centrosomes are inactivated, however, at later developmental stages (Stiess et al., 2010) and are dispensable for neuronal development in both mammalian and fly neurons (Nguyen et al., 2011; Stiess et al., 2010). No other active MTOCs within the neuronal soma have been described. Nevertheless, microtubules continue to grow within the soma (Nguyen et al., 2011; Sánchez-Huertas et al., 2016), and in mammalian neurons this depends in part on the HAUS complex (Sánchez-Huertas et al., 2016), which is also important for microtubule growth within axons and dendrites (Cunha-Ferreira et al., 2018; Sánchez-Huertas et al., 2016). Some MTOCs have been identified within dendrites: the basal body, or its surrounding region, within the distal part of *C. elegans* ciliated neurons acts as an MTOC, as does a similar region within the URX non-ciliated neuron (Harterink et al., 2018); an MTOC made from endosomes that tracks the dendritic growth cone in *C. elegans* PVD neurons has recently been identified (Liang et al., 2019); and fragments of Golgi called Golgi outposts within the dendrites of *Drosophila* dendritic arborisation (da) neurons are thought to recruit γ-TuRCs and act as MTOCs (Ori-McKenney et al., 2012; Yalgin et al., 2015; Zhou et al., 2014).

*Drosophila* larval da neurons are a popular *in vivo* neuronal model (Jan and Jan, 2010). Ease of imaging and genetic manipulation coupled with the ability to examine different neuronal classes, each with a stereotypical dendritic branching pattern, make them a model of choice when examining microtubule organisation within neurons. Four classes exist, including class I neurons that are proprioceptive and have the simplest “comb-like” dendritic branching pattern, and class IV neurons that are nociceptive and have the most elaborate branching pattern, tiling the surface of the larva (Figure S1) (Grueber et al., 2002). In both neuronal classes, microtubules in axons are predominantly plus-end-out throughout development, but microtubule polarity in dendrites progressively becomes more minus-end-out as the neurons develop; at two days post larval hatching the majority of dynamic microtubules are minus-end-out (Hill et al., 2012; Stone et al., 2008). Microtubule polarity within da neurons can be disrupted when microtubule regulators are depleted (Mattie et al., 2010; Nguyen et al., 2014; Ori-McKenney et al., 2012; Rolls and Jegla, 2015; Sears and Broihier, 2016; Weiner et al., 2016; Yalgin et al., 2015; Zhou et al., 2014), but we lack a full understanding of how microtubule polarity is established and maintained.

Golgi outposts are thought to provide localised sites of microtubule nucleation within the dendrites of da neurons to help regulate dendritic outgrowth and microtubule polarity (Ori-McKenney et al., 2012; Yalgin et al., 2015; Zhou et al., 2014). Unlike the somatic Golgi, which comprises cis, medial, and trans compartments organised into stacks and ribbons (Kondylis and Rabouille, 2009; Nakamura et al., 1995), Golgi outposts can be either single- or multi-compartment units with multi-compartment units having a higher propensity to initiate microtubule growth (Zhou et al., 2014). Within class I da neurons, Golgi outpost-mediated nucleation is thought to be dependent on the γ-TuRC-tethering protein Cnn and to help restrict dendritic branching (Yalgin et al., 2015), while within class IV neurons Golgi outpost-mediated nucleation is believed to be dependent on Pericentrin-like protein (Plp) and to help promote dendritic branching (Ori-McKenney et al., 2012). There remains some uncertainty, however, about the role of Golgi outposts in microtubule nucleation, as a separate study that examined the localisation of ectopically expressed γ-tubulin-GFP concluded that γ-tubulin localises predominantly to dendritic branch points in a Golgi outpost-independent manner (Nguyen et al., 2014).

In this study, we initially aimed to identify sites of microtubule nucleation within *Drosophila* da neurons. We used endogenously tagged γ-tubulin-GFP as a proxy for γ-TuRCs and performed a detailed analysis of γ-tubulin localisation within both class I and class IV da neurons. We find variation between neuronal classes in how γ-tubulin localise within dendrites, but find that the most prominent localisation of γ-tubulin is at the cis-compartment of somatic Golgi stacks within all sensory neurons of the dorsal cluster. This asymmetric γ-tubulin-Golgi association is not dependent on either Cnn or Plp, suggesting another molecule must regulate γ-TuRC recruitment to the Golgi.

Tracking of EB1-GFP comets within the soma suggests that microtubules are nucleated asymmetrically from the somatic Golgi. We find that these Golgi-derived microtubules initially grow preferentially towards the axon. During growth, they are also guided towards the axon and away from dendrites in a Kinesin-2-dependent manner, and they readily enter the axon contributing to plus-end-out microtubule polarity. In contrast, the relatively small number of microtubules that do grow towards a dendrite are normally excluded and this also depends upon Kinesin-2. After Kinesin-2 depletion, growing microtubules more readily approach and enter the dendrites causing a dramatic increase in the proportion of anterograde microtubules in proximal dendrites. This results in a reversal of overall microtubule polarity in the proximal dendrites from a predominantly minus-end-out microtubule population to a population where the majority of microtubules are plus-end-out. We therefore propose that plus-end-associated Kinesin-2 guides growing microtubules towards the axon and away from dendrites along a polarised microtubule network within the soma, while at dendrite entry points they generate a stalling force on growing microtubules when they encounter dendritic microtubules of opposite polarity.

## Results

### A detailed analysis of endogenous γ-tubulin localisation within class I and class IV da neurons

We began by examining the localisation of γ-tubulin (as a proxy for γ-TuRCs) within class I and class IV da neurons. To avoid any potential artefacts induced by ectopic overexpression, we used alleles where γ-tubulin23C (the zygotic form of γ-tubulin) was tagged at its endogenous locus with GFP ((Tovey et al., 2018) and this study). We generated fly stocks expressing two genetic copies of endogenously-tagged γ- tubulin23C-GFP (hereafter γ-tubulin-GFP) and the membrane marker mCD8-RFP, expressed either in class I or class IV da neurons, and imaged living animals. The most striking and obvious localisation of γ-tubulin-GFP was as multiple bright and relatively large puncta within the soma of both neuronal types (Figure 1A; Figure 2A); we address this localisation in subsequent sections. We could also detect discrete accumulations of γ-tubulin-GFP at specific dendritic sites (Figure 1A; Figure 2A), which were typically dim and varied between class I and class IV neurons. We therefore describe the localisation of γ-tubulin-GFP within dendrites for each neuron type in turn below.

**Figure 1.**
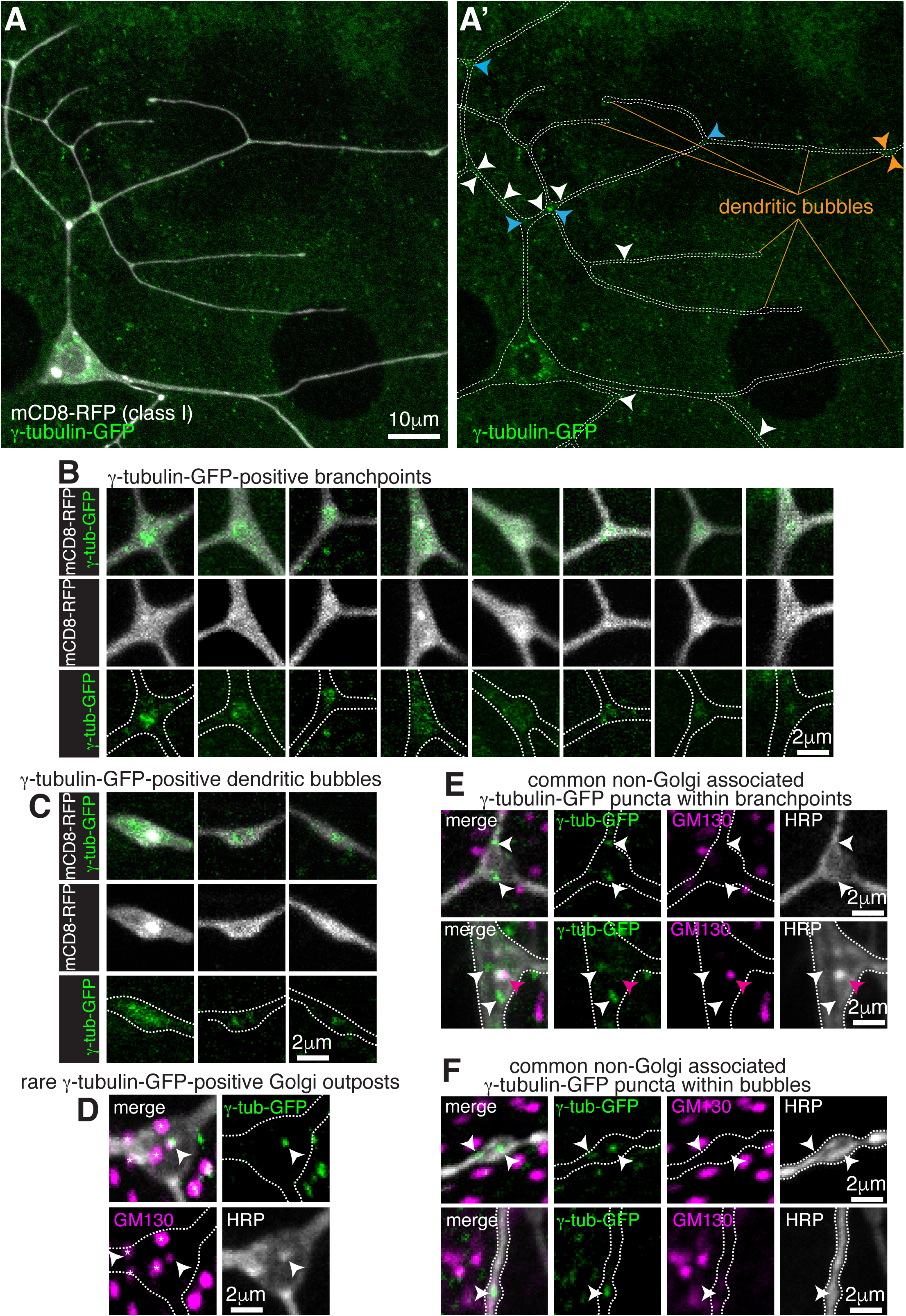
Endogenously-tagged γ-tubulin-GFP localises to a fraction of branchpoints and dendritic bubbles within class I da neurons. (**A**) Fluorescent confocal images of the proximal region of a class I da neuron expressing mCD8-RFP (greyscale) within a living 3^rd^ instar larva expressing endogenously-tagged γ-tubulin-GFP (green). Left panel (A) shows an overlay of the GFP and RFP signals, right panel (A’) shows only the GFP signal with the outline of the neuron drawn for clarity; white, blue and orange arrowheads indicate γ-tubulin-GFP puncta/accumulations within dendritic stretches, branchpoints, and dendritic bubbles, respectively. (**B,C**) Selected images of γ-tubulin-GFP-positive branchpoints (B) or dendritic bubbles (C) from living neurons as in (A). Individual mCD8-RFP channel images (greyscale) have been included for clarity. (**D- F**) Confocal images show branchpoints (D,E) or dendritic bubbles (F) from 3^rd^ instar larvae expressing endogenous γ-tubulin-GFP fixed and immunostained for GFP (green), GM130 (magenta) and HRP (greyscale). γ-tubulin-GFP signal was rarely observed co-localising with GM130 and HRP signal (D), and frequently observed independent of the Golgi markers at both branchpoints (E) and dendritic bubbles (F). See also Figure S1.

**Figure 2.**
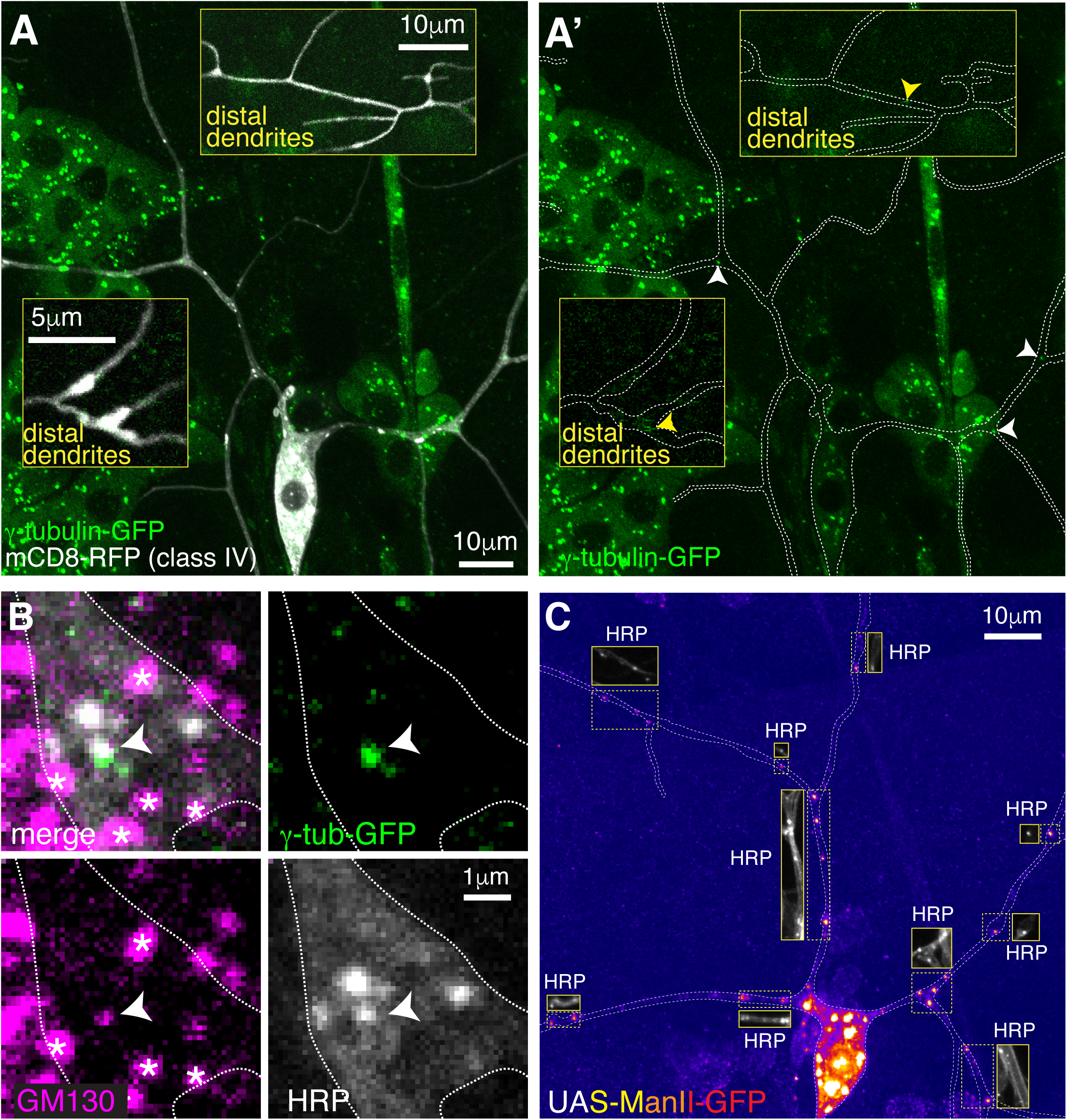
Endogenously-tagged γ-tubulin-GFP localises to a small fraction of proximal Golgi outposts within class IV da neurons. (**A**) Fluorescent confocal images of the proximal region of a class IV da neuron expressing UAS-mCD8-RFP (greyscale) within a living 3^rd^ instar larva expressing endogenously-tagged γ-tubulin-GFP (green). Left panel (A) shows an overlay of the GFP and RFP signals, right panel (A’) shows only the GFP signal with the outline of the neuron drawn for clarity; insets show examples of distal dendrites located >200μm from the soma; white and yellow arrowheads indicate bright or weak γ-tubulin-GFP puncta, respectively, within proximal or distal dendrites. (**B**) Images show a γ-tubulin-GFP-positive proximal branchpoint from a 3^rd^ instar larva expressing endogenous γ-tubulin-GFP fixed and immunostained for GFP (green), GM130 (magenta) and HRP (greyscale). Asterisks indicate GM130 foci from overlapping epidermal cells. Arrowhead indicates an example of a γ-tubulin-GFP-positive Golgi outpost observed within proximal class IV branchpoints. (**C**) Confocal images show the proximal region of a class IV dendritic arbor within a 3^rd^ instar larva expressing ManII-GFP (to mark medial Golgi) fixed and immunostained for GFP (fire) and HRP (greyscale within insets). Insets show HRP staining alone. See also Figure S1.

For class I da neurons we focussed on the ddaE neuron (hereafter class I neurons), as these have been the most widely characterised. From 13 class I neurons analysed, we detected an average of 1.1 discrete γ-tubulin-GFP puncta per 100μm of dendrite (Figure 1A). Most puncta were just above background intensity levels, although some were bright, and they were more frequently found away from branchpoints (80.2% of puncta) than within branchpoints (19.8% of puncta). We could detect γ-tubulin-GFP signal in 18% of branchpoints; in approximately half of these branchpoints the γ- tubulin-GFP appeared as puncta, often with multiple puncta per branchpoint, while in the other half of branchpoints the γ-tubulin-GFP signal was spread more diffusely through the branchpoint (Figure 1B). This diffuse signal was similar to that observed when γ-tubulin-GFP is overexpressed (Nguyen et al., 2014; Weiner et al., 2020) (Figure S1B), although it appeared that the frequency of branchpoint occupancy and the strength of the endogenous γ-tubulin-GFP signal was lower than that observed with ectopically expressed UAS-γ-tubulin-GFP. Nevertheless, this dendritic localisation that we observe likely represents the recently reported recruitment of γ- tubulin to endosomes at branchpoints (Weiner et al., 2020). We noticed that class I neurons display “bubbles”, where the diameter of the dendrite is locally increased to differing extents (Figure 1A); there were on average 2.6 “bubbles” per 100μm of dendrite with a larger fraction found further from the soma; ∼16.5% of bubbles contained γ-tubulin-GFP signal, either as weak puncta or a diffuse signal (Figure 1C), and ∼36.7% of the γ-tubulin-GFP puncta that we observed within dendrites were found within bubbles. To test whether γ-tubulin-GFP puncta within class I dendrites associate with Golgi outposts, we fixed and stained larval preparations expressing γ- tubulin-GFP with antibodies against HRP, which stain neuronal membranes, GFP, and the Golgi protein GM130. Out of 9 neurons from 3 larvae, we could only find one example where γ-tubulin-GFP colocalised with HRP staining, and in this case GM130 was also colocalised suggesting it was a Golgi outpost (Figure 1D). GM130 labels only multi-compartment Golgi outposts (Zhou et al., 2014) but HRP probably labels most, if not all, Golgi outposts. We chose not to use the ectopically expressed Golgi marker UAS-ManII-GFP used in some previous studies, as this has been reported to “leak” into endosomes within the dendrites of class I da neurons (Weiner et al., 2020). Consistent with this, we found that UAS-ManII-GFP can form puncta, enrichments, and long stretches within class I dendrites that are not associated with HRP staining (Figure S1C). These apparent cytosolic accumulations may therefore be due to an excess of ManII protein within the dendrites. This appeared to be specific to dendrites, however, as UAS-ManII-GFP always colocalised with HRP staining within the soma (Figure S1D), possibly due to the higher concentration of Golgi within the soma. All γ- tubulin-GFP puncta (except for the single occasion noted above), including those within branchpoints, dendrites and bubbles, did not colocalise with HRP or GM130 staining (Figure 1E,F). Thus, our data strongly suggests that γ-TuRCs do not typically associate with Golgi outposts within class I neurons, and instead localise to a fraction of branchpoints and dendritic bubbles in a Golgi outpost-independent manner.

For class IV neurons we focussed on the ddaC neuron (hereafter class IV neurons), as these have also been the most widely characterised. We analysed two distinct dendritic regions that we term proximal (within ∼100μm of the soma) and distal (over ∼200μm from the soma). We detected an average of 2.3 and 0.3 γ-tubulin-GFP puncta per 100μm of dendrite within proximal (10 neurons analysed) and distal (7 neurons analysed) regions, respectively, and the intensity of the majority was only just above background levels, including all of the puncta in the distal regions (Figure 2A). Nevertheless, we consistently observed bright γ-tubulin-GFP puncta within the primary and secondary branchpoints close to the soma (Figure 2A). 44.6% of γ-tubulin-GFP puncta observed in the proximal regions were found within the large primary and secondary branchpoints, and 57% of these branchpoints contained γ-tubulin-GFP puncta, which were always bright relative to other foci within the dendrites (Figure 2A). Staining with HRP and GM130 antibodies showed that these bright γ-tubulin-GFP puncta associated with Golgi outposts (Figure 2B). Intriguingly, γ-tubulin-GFP only colocalised with HRP puncta that also associated with GM130 (Figure 2B), suggesting that γ-TuRCs are recruited only to multi-compartment Golgi outposts. The high frequency of proximal branchpoints that contained γ-tubulin-GFP puncta contrasted with the near absence of γ-tubulin-GFP puncta within the smaller branchpoints of the distal arbor, where only 1.7% of branchpoints contained detectable but very weak γ- tubulin-GFP signal (Figure 2A insets). In contrast to class I neurons, we found that the vast majority of ectopically expressed ManII-GFP puncta within class IV neurons colocalised with HRP and did not form large accumulations or stretches within the dendrites (Figure 2C), suggesting that ectopically expressed ManII-GFP can be used as a reliable marker of Golgi outposts in class IV neurons. We observed an average of ∼12 and ∼5 ManII-GFP-positive Golgi outposts within proximal and distal regions, respectively, which is consistent with previous observations (Zheng et al., 2008) and far higher than the 2.3 and 0.3 γ-tubulin-GFP puncta per 100μm that we observed in proximal and distal dendrites (compare Figure 2A to Figure 2C). Thus, only Golgi outposts within the proximal branchpoints of class IV neurons associate readily with γ-tubulin. In contrast to class I neurons, we very rarely observed γ-tubulin-GFP spread diffusely through branchpoints, suggesting that Golgi outpost-independent accumulation of γ-tubulin at branchpoints is specific to class I neurons. Moreover, we found far fewer dendritic bubbles per 100μm of dendrite in class IV neurons (0.4 and 0.8 per 100μm dendrite in proximal and distal regions, respectively). Collectively, our data show that Golgi outposts within class IV neuron branchpoints close to the soma frequently associate with γ-tubulin, but that the majority of Golgi outposts do not. Whether this proximal Golgi outpost associated γ-tubulin represents fully functional γ- TuRCs remains to be tested.

### γ-tubulin-GFP localises to the somatic Golgi of sensory neurons in a Cnn- and Plp-independent manner

The most striking and obvious localisation of γ-tubulin-GFP within both class I and class IV neurons was as multiple large bright puncta within their soma (see images of neurons from live animals in Figure 1A and Figure 2A, and of fixed and stained neurons in Figure 3A-C). We also observed similar puncta within the somas of other nearby sensory neurons (Figure 3A), including external sensory (es) neurons; these sensory neurons possess basal bodies that appear to associate with large amounts of γ-tubulin-GFP (Figure 3A). Staining with antibodies against HRP and the Golgi marker GM130 showed that all γ-tubulin-GFP puncta within the soma of all sensory neurons associated with somatic Golgi stacks (Figure 3B,C; Figure S2), which are typically scattered throughout the cytosol in *Drosophila* cells while still maintaining cis-medial-trans polarity (Kondylis and Rabouille, 2009). These Golgi stacks can be oriented side-on or face-on to the imaging plane, appearing either more elongated or circular, respectively. Our staining showed that the signals of γ-tubulin-GFP and GM130 partially overlapped at side-on stacks, with γ-tubulin-GFP extending further out laterally than GM130 (Figure 3B; Figure S2A); γ-tubulin-GFP surrounded GM130 in a ring-like pattern on face-on stacks (Figure 3C; Figure S2B). To determine whether γ- tubulin associates specifically with the cis-Golgi, as has been suggested for γ-tubulin in non-neuronal mammalian cells (Wu et al., 2016), we stained the neurons with antibodies against HRP, GM130, and Arl1. The mammalian homologue of GM130 is a cis-Golgi protein, while Arl1 is a known trans Golgi protein in *Drosophila* (Munro, 2011). We found that the GM130 and Arl1 signals were offset at side-on stacks, consistent with them being cis- and trans-Golgi proteins, respectively. Moreover, γ- tubulin colocalised with GM130 rather than Arl1 (Figure 3D). Together, this shows that γ-tubulin-GFP, possibly in the form of γ-TuRCs, localises to the rims of the cis-Golgi stack in the soma of *Drosophila* da neurons.

**Figure 3.**
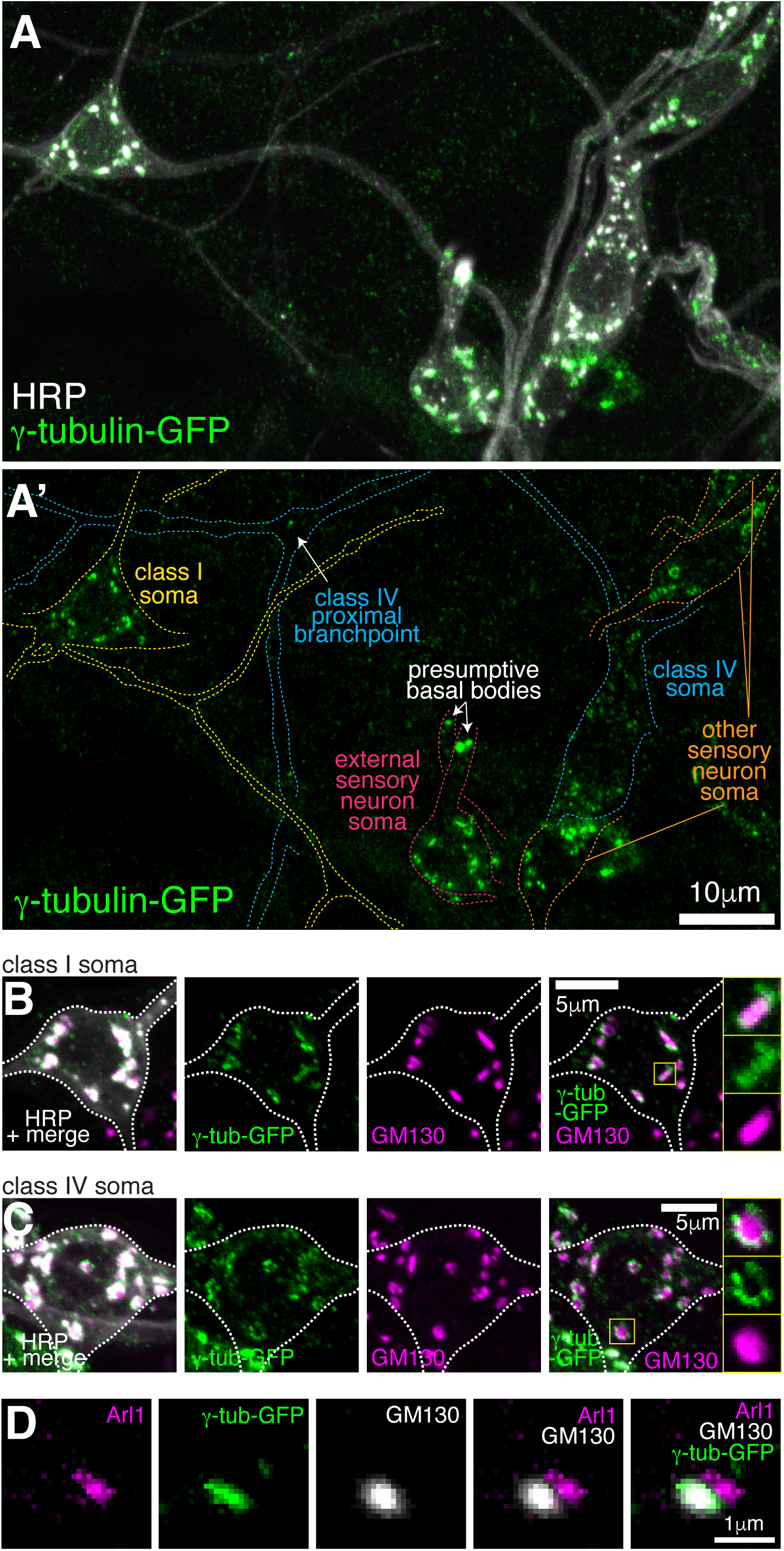
Endogenously-tagged γ-tubulin-GFP localises to the somatic Golgi of sensory neurons. (**A**) Confocal images show the somas and some proximal dendrites of sensory neurons within the dorsal cluster: class I (yellow), class IV (blue), as (pink) and other (orange) from a 3^rd^ instar larva expressing endogenously-tagged γ-tubulin-GFP and immunostained for GFP (green) and HRP (marking Golgi stacks, greyscale). Upper panel (A) shows an overlay of the GFP and HRP signals, lower panel (A’) shows only the GFP signal with coloured neuronal outlines drawn for clarity. (**B, C**) Confocal images show the somas of class I (B) or class IV (C) da neurons from a 3^rd^ instar larva expressing endogenously-tagged γ-tubulin-GFP fixed and immunostained for GFP (green), GM130 (magenta) and HRP (greyscale). Enlarged boxes in (B) and (C) show side-on and face-on stacks, respectively. (**D**) Confocal images show an example of a single Golgi stack within a da neuron from a 3^rd^ instar larva expressing endogenously-tagged γ-tubulin-GFP fixed and immunostained for GFP (green), Arl1 (trans-Golgi, magenta) and GM130 (cis-Golgi, greyscale). The larva was also stained with HRP antibodies to identify neurons, but this channel has been omitted for clarity. See also Figure S2.

Two proteins, Cnn and Plp, were possible candidates for γ-TuRC recruitment to the somatic Golgi, as their homologues are required for proper organisation of microtubules at the somatic Golgi in cycling mammalian cells (Rios, 2014; Roubin et al., 2013; Wang et al., 2010; Wu et al., 2016), and Cnn and Plp have been implicated in γ-TuRC recruitment to Golgi outposts in *Drosophila* class I and class IV neurons, respectively (Ori-McKenney et al., 2012; Yalgin et al., 2015). Cnn is a multi-isoform gene with three sets of isoforms driven by three different promoters (Eisman et al., 2009) (Figure S3A). Promoter one drives the best-studied isoform (that we term Cnn-P1) that localises to centrosomes during mitosis; promoter two drives isoforms (Cnn-P2) that are yet to be characterised; and promoter three drives isoforms that are expressed specifically within testes (Chen et al., 2017) and so have not been considered in this study. Immunostaining with antibodies against Cnn revealed very weak, if any, signal within the soma or dendrites of the dorsal sensory neurons; while the presumptive basal bodies of the es neurons displayed a strong Cnn signal (data not shown). Given that antibody staining can be problematic, we generated flies where Cnn-P1 or Cnn-P2 were tagged with GFP at their isoform-specific N-termini (hereafter, GFP-Cnn-P1 and GFP-Cnn-P2; Figure S3A). The GFP insertions appear functional as flies could be kept as homozygous stocks and the localisation of GFP-Cnn-P1 to centrosomes in syncytial embryos was normal (Movie S1). GFP-Cnn-P1 signal was very weak and inconsistent within the soma of living class I da neurons (Figure 4A). In contrast, we could readily detect clear GFP-Cnn-P1 puncta within distal dendritic bubbles in live animals (Figure 4B). In fixed samples, there was a weak GFP-Cnn-P1 signal associated with the somatic Golgi in the dorsal cluster of sensory neurons, similar to that observed in live samples (Figure S3B), although we did not detect GFP-Cnn-P1 associated with HRP puncta within dendrites (Figure S3B; data not shown). We found a strong GFP-Cnn-P1 signal at the presumptive basal bodies of the es neurons (Figure S3B). Collectively our data suggest that GFP-Cnn-P1 does not readily associate with Golgi, but accumulates within a fraction of dendritic bubbles in class I neurons, possibly together with γ-tubulin.

**Figure 4.**
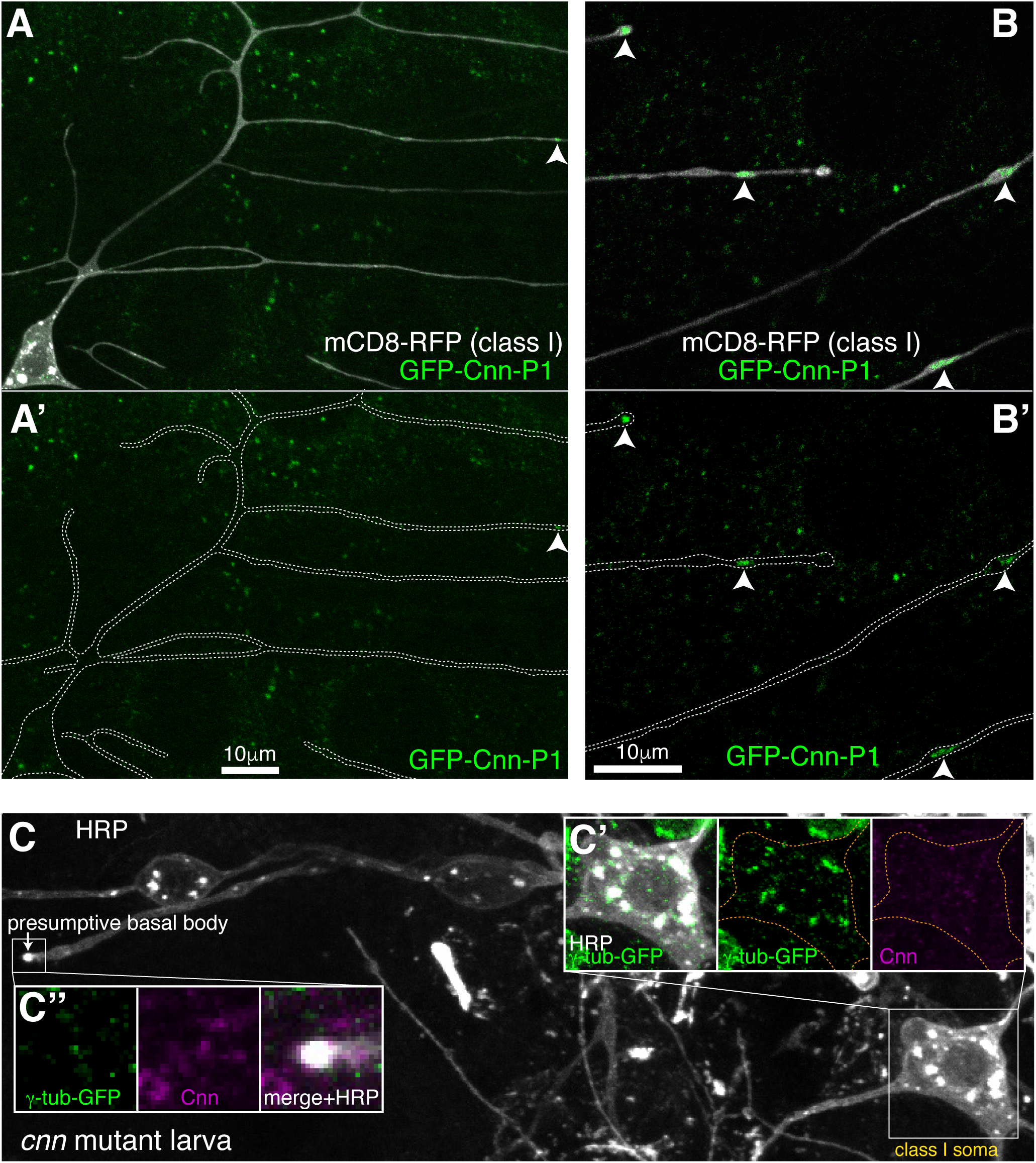
Cnn-P1 is dispensable for γ-tubulin-GFP recruitment to the somatic Golgi. (**A,B**) Fluorescent confocal images of proximal (A) and distal (B) regions of class I da neurons expressing mCD8-RFP (greyscale) within a living 3^rd^ instar larva expressing endogenously-tagged GFP-Cnn-P1 (green). Upper panels show an overlay of the GFP and RFP channels, lower panels show only the GFP channel with the outline of the neurons drawn for clarity. Arrowhead in (A) indicates a rare dendritic GFP-Cnn-P1 puncta within a proximal dendritic bubble, while arrowheads in (B) indicate more frequent GFP-Cnn-P1 accumulations within distal dendritic bubbles. (**C**) Confocal images show the somas and some proximal dendrites of sensory neurons within the dorsal cluster from a 3^rd^ instar *cnn* mutant larva expressing endogenously-tagged γ- tubulin-GFP and immunostained for GFP (green), HRP (greyscale) and Cnn (magenta). Images in (C’) and (C’’) show the separate channels for the class I soma and the presumptive basal body of an es neuron, respectively. See also Figure S3.

In contrast to GFP-Cnn-P1, we did not detect any obvious GFP-Cnn-P2 signal within the soma or dendrites of da neurons in living larvae (Figure S3C). Instead, GFP-Cnn-P2 appeared to be expressed within glial cells that ensheath the axons, soma, and part of the proximal dendritic region of the da neurons (Figure S3C). These glia are known to help regulate dendritic development (Han et al., 2011; Sepp and Auld, 2003; Yadav et al., 2019), and thus Cnn-P2 isoforms may have an indirect role in dendritic arbor growth. GFP-Cnn-P2 also localised strongly to the presumptive basal body of es neurons (data not shown). Consistent with the localisation pattern of GFP-Cnn-P1 and GFP-Cnn-P2, γ-tubulin-GFP could still associate with the somatic Golgi of the sensory neurons in *cnn* mutant larvae, but not to the basal bodies of es neurons (Figure 4C). We therefore conclude that Cnn is dispensable for γ-TuRC recruitment to the somatic Golgi within sensory neurons.

In contrast to Cnn, antibodies against Plp readily stained the somatic Golgi in all sensory neurons, including class I and class IV da neurons (Figure S4A). The Plp signal was, however, offset from the γ-tubulin-GFP signal at Golgi stacks in both class I and class IV da neurons (Figure 5A,B), suggesting that they localise to different Golgi compartments. Plp also associated with the class IV proximal Golgi outposts (Figure 5C) and the presumptive basal bodies of the es neurons (Figure S4B), and was enriched within a fraction of the distal class I dendritic bubbles (Figure S4C). Strikingly, γ-tubulin-GFP localisation at the somatic Golgi of all sensory neurons, including class I and class IV neurons was unaffected in *plp* mutant larvae (Figure 5D). This was also true of the γ-tubulin-positive proximal Golgi outposts in class IV neurons (Figure 5D). The absence of Plp did, however, lead to the loss of γ-tubulin-GFP from the basal bodies of es neurons (Figure 5D). In summary, neither Plp nor Cnn are required for the efficient recruitment of γ-tubulin to the somatic Golgi within sensory neurons, or to the few γ-tubulin-GFP-positive Golgi outposts within the proximal branch points of class IV da neurons, but both are required for the localisation of γ-tubulin-GFP to the basal body region within es neurons.

**Figure 5.**
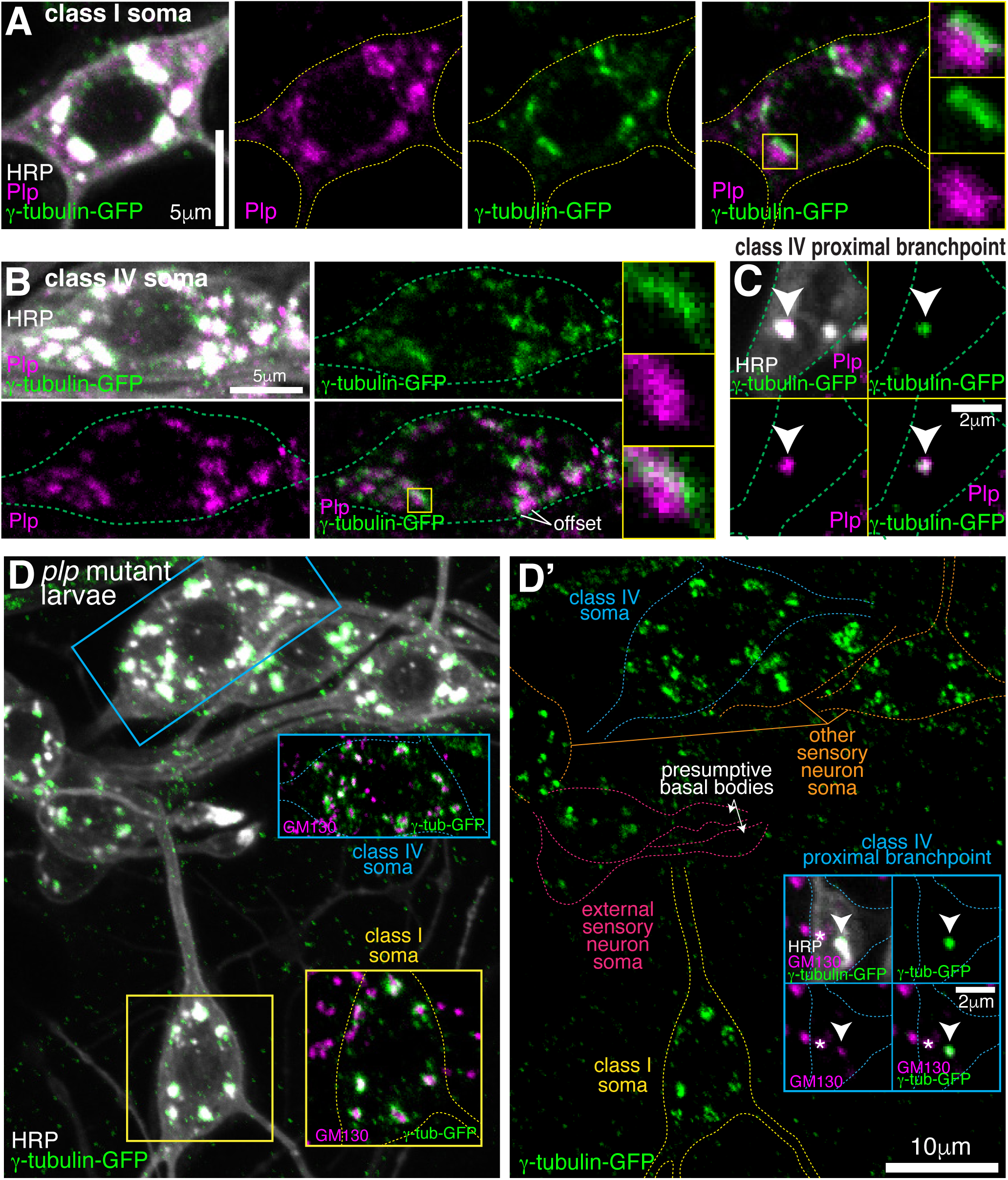
Endogenous Plp localises to Golgi but is dispensable for γ-tubulin-GFP recruitment. (**A-C**) Confocal images show the somas of a class I (A) or a class IV (B) neuron and a proximal branchpoint of a class IV neuron (C) from 3^rd^ instar larvae expressing γ-tubulin-GFP fixed and immunostained for GFP (green), Plp (magenta) and HRP (greyscale). Arrowhead in (C) indicates a γ-tubulin-GFP-positive Golgi outpost that contains Plp. (**D**) Confocal images show the somas and some proximal dendrites of sensory neurons within the dorsal cluster: class I (yellow), class IV (blue), es (pink) and other (orange), from a *plp* mutant 3^rd^ instar larva expressing endogenously-tagged γ-tubulin-GFP and immunostained for GFP (green), HRP (greyscale) and GM130 (magenta). Left panel (D) shows an overlay of the GFP and HRP signal, with insets showing overlays for different neurons, as indicated; right panel (D’) shows the GFP channel with coloured outlines of the different neurons drawn for clarity, with insets showing separate channels for a proximal branchpoint. See also Figure S4.

### Growing microtubules originate asymmetrically from the somatic Golgi

We next wanted to assess whether the somatic Golgi is an active MTOC. EB1-GFP binds to growing microtubule ends and generates fluorescent “comets”, typically representing growing microtubule plus ends. While the origin of these comets can represent either microtubule regrowth after catastrophe or sites of microtubule nucleation, the position of emerging EB1-GFP comets has been routinely used in *Drosophila* neurons as a proxy for nucleation sites (Nguyen et al., 2014; Ori-McKenney et al., 2012; Weiner et al., 2020; Yalgin et al., 2015; Zhou et al., 2014). It is generally considered that comets repeatedly emerging from the same location are likely to represent nucleation sites (Ori-McKenney et al., 2012; Zhou et al., 2014). We therefore imaged the soma of class I neurons expressing EB1-GFP and the Golgi marker ManII-mCherry and manually tracked each EB1-GFP comet (Figure 6A; Movie S2; the circle of each track represents the latest position of the EB1 comet). Comets could be observed originating at both Golgi stacks and non-Golgi sites. They emerged at different Golgi stacks within the same cell, and sequentially from the same Golgi stack, suggestive of nucleation. Intriguingly, many Golgi-derived comets appeared to grow towards the axon. Measuring the initial angle of each comet’s growth relative to the axon entry site (indicated in Figure 6A) showed that there was a strong bias for initial growth towards the axon (163 comets analysed from 59 Golgi stacks from 7 neurons, p<0.001), although some comets did grow away from the axon and thus towards dendrites (Figure 6B). While it is possible that the direction of comet growth can be influenced by comets growing into the soma from the dendrites, which will tend to grow towards the axon, we minimised this effect by quantifying only those comets that originate from Golgi stacks. The comets we analysed are therefore more likely to represent true growth events from the Golgi rather than catastrophe and regrowth of microtubules that originated from dendrites.

**Figure 6.**
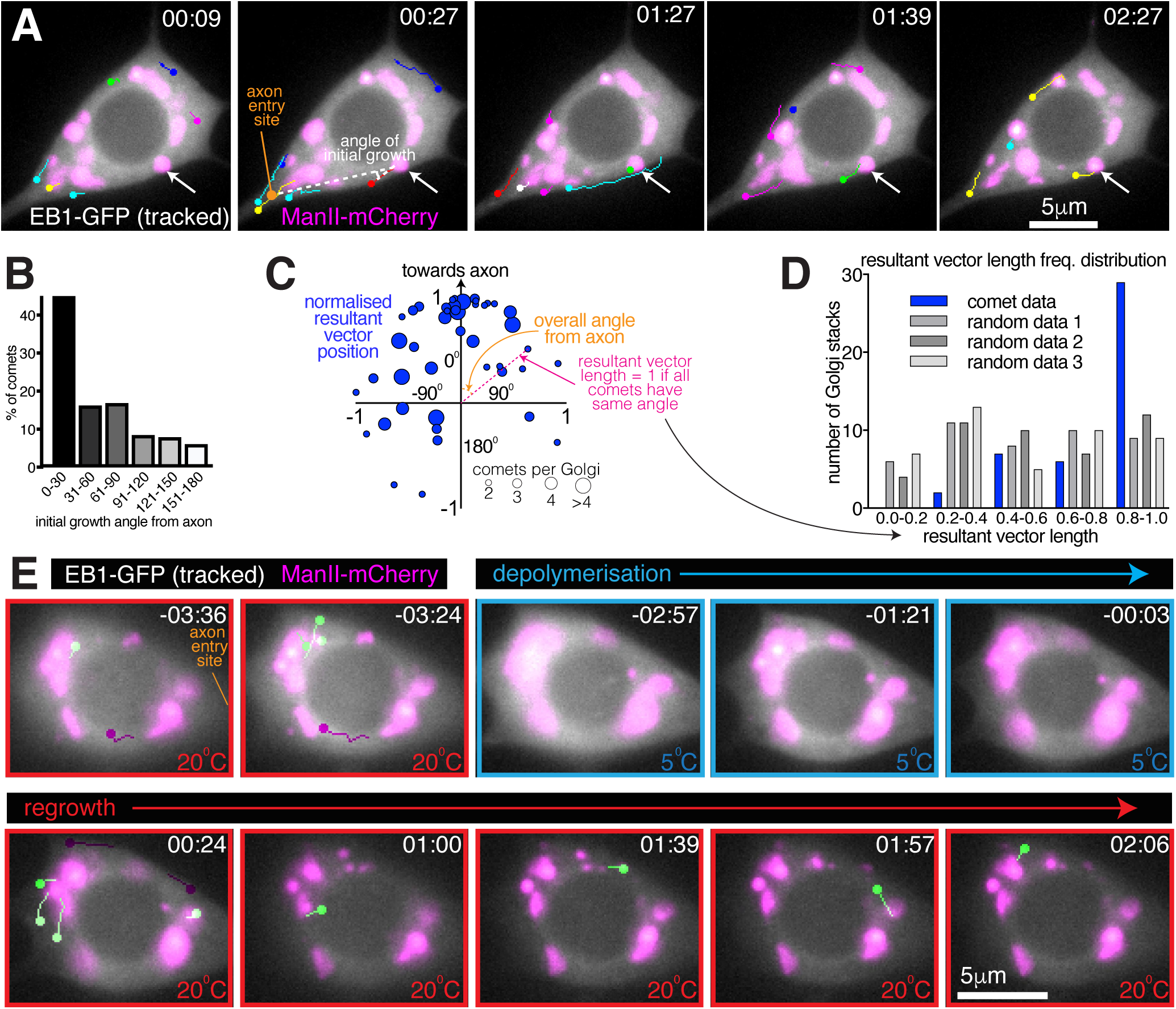
Somatic Golgi stacks nucleate microtubules. (**A**) Widefield fluorescent images from a movie showing the soma of a class I da neuron expressing EB1-GFP (greyscale) and ManII-mCherry (magenta). Manually assigned multi-colour tracks of EB1-GFP comets were drawn over each image (filled circle = last location of comet). Arrow indicates a Golgi stack from which sequential EB1-GFP comets emerge; dotted lines show an example of an angle measurement of a comet’s initial growth relative to the axon entry point (orange circle). Time in min:s from the start of the movie is shown. (**B**) Graph shows a frequency distribution of the initial growth angle (before turning) relative to the axon entry site (as indicated in (A)) of EB1-GFP comets that emerged from Golgi stacks. Negative angles were made positive so as not to distinguish between comets growing to the left or right of the axon. 163 comets from 59 Golgi stacks from 7 neurons were analysed. (**C**) Scatter graph shows the position of the normalised resultant comet vectors (see methods) for each of 44 Golgi stacks from 7 neurons. Circle position is representative of the average angle from the axon and how similar the angles were (as indicated). Circle size reflects the number of comets analysed per Golgi stack (as indicated below). (**D**) Graph shows a frequency distribution of the resultant vector lengths for the comet data (blue) and for 3 datasets produced using randomly generated angles (grey shades). (**E**) Widefield fluorescent images from a movie showing the soma of class I da neuron expressing UAS-EB1- GFP (greyscale) and UAS-ManII-mCherry (magenta). The sample was subjected to a warm-cool-warm temperature cycle to induce depolymerisation (5°C; blue outline) and then repolymerisation (20°C; red outline) of dynamic microtubules. Tracks were manually colour coded: green = comets emerging from Golgi stacks; purple = comets emerging from elsewhere. Time in min:s from the 5°C to 20°C transition is shown. See also Movie S2 and Movie S3.

We also noticed that the direction of initial growth of each comet that emerged from the same Golgi stack was similar, irrespective of their angle from the axon. We therefore generated and plotted normalised resultant vectors (final vector position from a (0,0) origin represented by a blue circle) for the comets from each Golgi stack that produced two or more comets (Figure 6C; see Methods). The angle between the positive Y-axis and a line connecting (0,0) and a given circle on the graph is representative of the overall comet angle from the axon; the distance (d) of each circle from (0,0) is representative of the similarity of comet angles (d=1 if all angles are the same; d=0 if all angles are evenly distributed). 77% of the circles were in the upper quadrants (Figure 6C; p<0.001), again showing that there was a preference for comets to grow initially towards the axon. Moreover, there was a bias for resultant vector lengths to be large, as compared to randomly generated data (Figure 6D; p<0.001). We conclude that comets from a particular Golgi stack emerge within a small angle with respect to each other and with a directional preference towards the axon. While not definitive, the similarity in the direction of comets emerging repeatedly from the same Golgi stack is indicative of consecutive microtubule nucleation events.

### A Temperature-based microtubule nucleation assay suggests that microtubules are nucleated from the somatic Golgi in class I da neurons

The origin of EB1-GFP comets is not a perfect proxy for microtubule nucleation, as EB1-GFP comets also appear during the regrowth of partially depolymerised microtubules. We therefore performed a microtubule nucleation assay using a temperature-control device (CherryTemp, Cherry Biotech) to cool samples rapidly to 5°C and then re-heat them to 20°C during continuous imaging of the sample. Cooling typically causes depolymerisation of dynamic microtubules and warming causes their regrowth. Cooling-warming microtubule nucleation assays have previously been performed in various systems, including *Drosophila* embryos (Hayward et al., 2014), *Drosophila* S2 cells (Bucciarelli et al., 2009), and in mammalian cells (Torosantucci et al., 2008). While populations of cold-stable microtubules have been identified in mammalian neurons, cold stability is thought to be induced by binding to MAP6 proteins, which are specific to vertebrates (Bosc et al., 2003; Delphin et al., 2012). We therefore expected that cooling would result in the depolymerisation of at least the dynamic microtubules within *Drosophila* neurons, allowing us to correlate the position of new comet growth with nucleation sites.

During cooling-warming cycles we observed no obvious effect on the distribution of the somatic Golgi stacks (Figure 6E; Movie S3), although thermal-fluctuation-induced movement of the glass coverslip made it difficult to follow the first few timepoints after cooling or warming. When the appropriate focal plane was reached shortly after warming, we could observe EB1-GFP comets emerging from the ManII-mCherry-labelled Golgi structures (green-labelled comets, Figure 6E; Movie S3), suggestive of Golgi-located microtubule nucleation events. Some comets emerged from non-Golgi sites (purple-labelled comets, Figure 6E; Movie S3), suggesting that other sites of microtubule nucleation may exist. This would not be surprising, given that HAUS, the mammalian homologue of *Drosophila* Augmin that recruits γ-TuRCs to the sides of pre-existing microtubules, is required for microtubule nucleation events within the soma of mammalian neurons in culture (Sánchez-Huertas et al., 2016). It may also be, however, that the non-Golgi comets grew initially from Golgi stacks in a different focal plane or that these comets represented the re-growth of microtubules that were not fully depolymerised. Indeed, while EB1-GFP comets disappeared rapidly on cooling, suggesting that dynamic microtubules quickly lose their GTP-tubulin cap and thus depolymerise, it is possible that a proportion of microtubules within the soma are cold stable and therefore won’t depolymerise. Presumably, most of these stable microtubules would remain stable, but it is possible that some would start to grow and thus contribute to the EB1-GFP comets that we observe on warming.

In our opinion, the best evidence for microtubule nucleation occurring at the somatic Golgi comes from the observation that several comets emerged from Golgi stacks relatively late after warming. For example, in Movie S3 four comets emerged from Golgi stacks at least 50s post warming. While some of these late comets emerged from a Golgi stack that had generated a comet immediately after warming, others emerged from Golgi stacks that had not generated a comet immediately after warming. Most importantly, the direction of all late emerging comets suggests that they were not simply generated by catastrophe-rescue of a microtubule that had grown immediately after warming. This strongly suggests that these late emerging comets represent genuine nucleation events from the Golgi, although it is impossible to rule out that they could have been generated by re-growing microtubules that were originally out of focus.

### Growing microtubules within the soma are guided towards the axon while being excluded from entering dendrites in a Kinesin-2-dependent manner

We next wanted to determine the fate of growing microtubules within the soma. We therefore imaged and tracked EB1-GFP comets within the soma and proximal axons and dendrites of 13 class I da neurons (Figure 7A; Movie S4). These neurons have at least two primary dendrites but only a single axon; however, comets that initiated within the soma often grew into the axon, while few entered dendrites (Movie S4). We found that this was due to two major factors. The first was that a higher proportion of comets reached the axon entry site: of the 666 comets that had initiated within the soma across all movies 104 approached the entrance to the axon (15.6%), while only 47 approached the entrance of a dendrite (7.1%) (Figure 7C; p<0.001); the remaining 77.3% of comets terminated within the soma away from the axon or dendrites. The second important factor was that when comets arrived at either the axon entry site or a dendritic entry site, they had more chance of entering the axon: of the 104 comets that approached the entrance to an axon across all movies 56 entered (53.8%), while of the 47 comets that approached the entrance to a dendrite only 13 entered (27.7%) (Figure 7D; p<0.001). Overall, 8.4% of all comets that originated in the soma entered the axon while only 2.0% entered a dendrite. In summary, growing microtubule plus ends within the soma preferentially reach the entrance to the axon and can readily enter, while the few that reach dendrites are normally excluded from entering.

**Figure 7.**
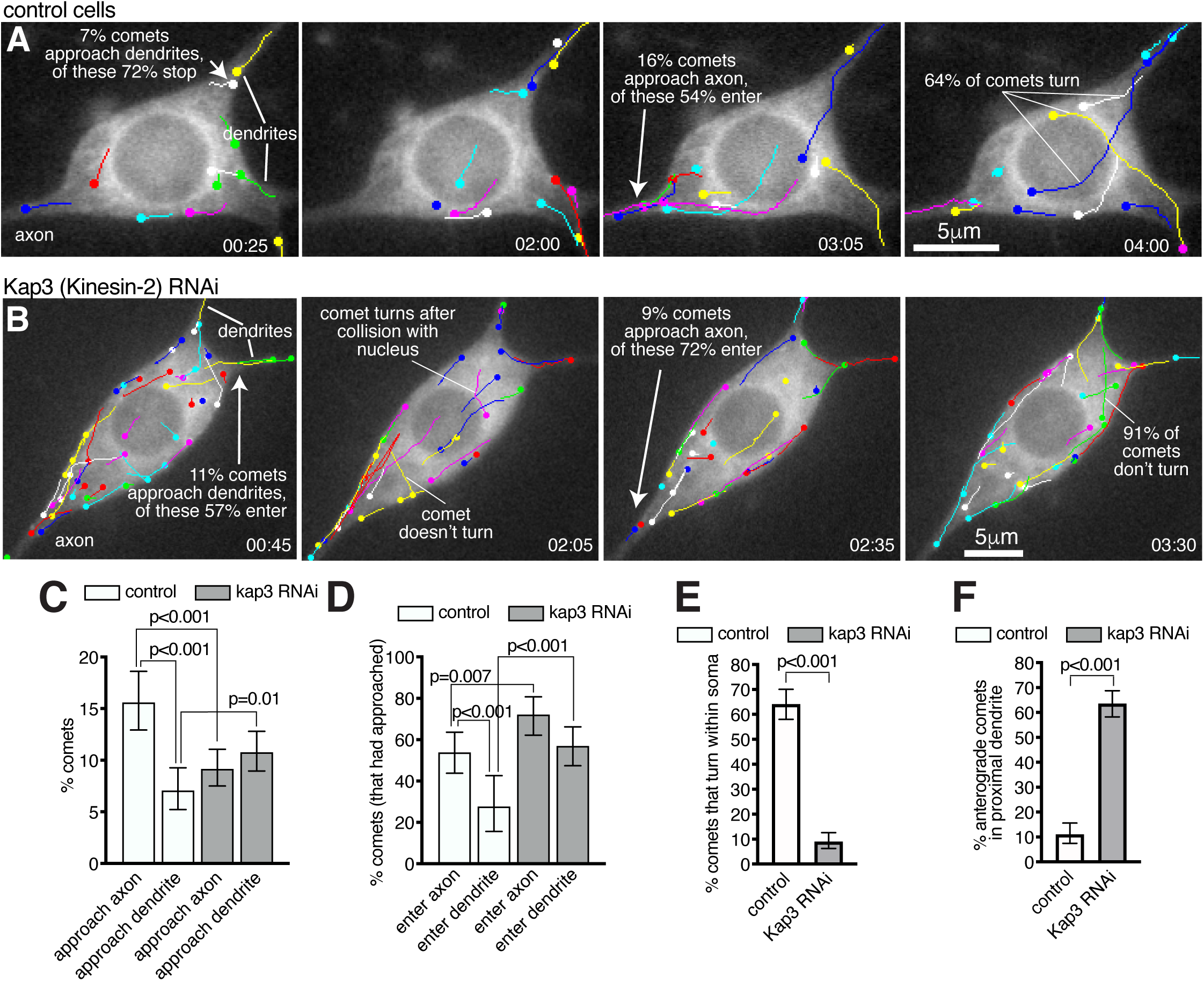
Microtubules within the soma grow preferentially towards the axon and are excluded from entering dendrites in a Kinesin-2-dependent manner. (**A,B**) Widefield fluorescent images from movies showing the somas of class I da neurons expressing UAS-EB1-GFP and either γ-tubulin-37C RNAi (control) (A) or UAS-Kap3 RNAi (B). Manually assigned multi-colour tracks of EB1-GFP comets are drawn over each image (filled circle = last location of comet). Time in min:s from the start of the movie is shown. (**C**) Graph shows the % of comets that approach either the axon (green) or the dendrites (magenta) in either control (666 comets analysed across 13 movies) or Kinesin-2 RNAi (1058 comets analysed across 9 movies) class I da neuron somas, as indicated. (**D**) Graph shows the % of comets that enter the axon (green) or the dendrites (magenta) as a proportion of those that had approached the axon in either control or Kinesin-2 RNAi class I da neuron somas, as indicated. (**E**) Graph shows the % of applicable comets that display turning events within either control (n=257 comets across 13 movies) or Kinesin-2 RNAi (n=386 comets across 9 movies) class I da neuron somas, as indicated. (**F**) Graph shows the % of comets that are anterograde in the proximal primary dendrite (before any branches) in either control (n=252 comets across 13 movies) or Kinesin-2 RNAi (n=338 comets across 9 movies) class I da neurons, as indicated. Error bars in (C-F) show the 95% confidence intervals.

While asymmetric nucleation from the somatic Golgi likely contributes to more microtubule plus ends reaching the axon, we also observed several occasions where microtubules turned towards the axon (Figure 7A; Movie S4). Microtubule turning events have been observed previously within dendritic branchpoints of da neurons, where the microtubules turn towards the soma along stable microtubules (Mattie et al., 2010; Weiner et al., 2016). This depends upon Kinesin-2, which is a heterotrimeric plus-end-directed motor whose regulatory subunit, Kap3, interacts with EB1 via APC (Mattie et al., 2010). It has therefore been proposed that plus-end-associated Kinesin-2 guides growing microtubules along and towards the plus end of so-called “rail” microtubules, something that can be recapitulated *in vitro* (Chen et al., 2014; Doodhi et al., 2014). When we depleted Kap3 from class I da neurons by RNAi there was a dramatic reduction in the frequency of microtubule turning events within the soma. In control cells, of the 257 comets across all movies that grew for more than 2μm within the soma and that did not travel along the cell cortex, 165 displayed turning behaviour (64.2%), while across 9 Kap3 RNAi cells only 35 of the 386 qualifying comets displayed turning behaviour (9.1%) (Figure 7A,B,E; Movie S4,5; p<0.001). Comets in Kap3 RNAi cells tended to change direction only when travelling along the cell cortex or after collision with the nuclear envelope or cell cortex (Figure 7B; Movie S5). As a result of reduced turning in Kap3 RNAi neurons, a lower proportion of comets reached the axon entry site and a higher proportion reached dendritic entry sites, compared to control cells: of the 1,058 comets that originated within the soma of Kap3 RNAi cells, 97 approached the axon entry site (9.2%, compared to 15.6% in control neurons, p<0.001) and 114 approached a dendritic entry site (10.8%, compared to 7.1% in control neurons, p=0.0098) (Figure 7C). These data suggest that Kinesin-2 guides growing plus ends along pre-existing microtubules towards the axon and away from dendrites within the soma.

Comets that did arrive at the axon entry site could still readily enter the axon after Kap3 RNAi (Figure 7D); there was actually a ∼1.3-fold increase compared to controls in the proportion of comets entering the axon (72.2% versus 53.5%, p=0.007). There is no requirement for Kinesin-2 for microtubule growth *per se*, which is presumably why microtubules can still enter once they reach the axon. It is unclear why entry is increased, but it is possible that Kinesin-2 may limit plus-end growth in general, although this purely speculative. Importantly, increased entry of growing microtubules into the axon has no major effect on microtubule polarity, as axons normally contain predominantly plus-end-out microtubules.

Most significantly, the proportion of comets that entered dendrites after Kap3 RNAi increased ∼2-fold compared to controls, a much higher increase than that seen for axons. Of the 114 comets across 9 movies that approached the entrance to a dendrite in Kap3 RNAi neurons, 65 entered (57.0%, compared to 27.7% in control neurons; p<0.001; Figure 7D). This affect was even more striking when knocking down a different Kinesin-2 component, Klp64D, where 56 of the 80 comets (across 10 movies) that approached a dendrite could enter (70.0%; p<0.001; Movie S6). Increased entry of comets into dendrites contributed to, if not caused, a dramatic increase in the proportion of anterograde comets within proximal primary dendrites (prior to any dendritic branches): in control neurons, the vast majority of comets that we could observe in all proximal dendrites (88.9%) were retrograde (minus-end-out microtubules), while in Kap3 RNAi neurons the majority (63.6%) were anterograde (plus-end-out microtubules) (Figure 7F; p<0.001). An explanation for Kinesin-2 being required to prevent growing microtubules from entering dendrites may be that these microtubules need to grow past dendritic microtubules of opposite polarity. If plus-end-associated Kinesin-2 engages with these oppositely polarised microtubules it would presumably create a backward stalling force when trying to ‘walk’ towards the plus end of the dendritic microtubule (see discussion for more detail).

We conclude that Kinesin-2 is required to guide growing microtubule plus ends within the soma towards the axon and to prevent them from entering dendrites, which is essential in order to maintain minus-end-out microtubule polarity within proximal dendrites.

### A model for microtubule regulation within the neuronal soma

Based on our observations, and the findings from previous studies (Chen et al., 2014; Doodhi et al., 2014; Mattie et al., 2010; Weiner et al., 2016), we propose a model to explain how microtubules are generated and regulated within the soma of da neurons, which also helps explain how microtubule polarity is promoted in axons and maintained within proximal dendrites (Figure 8). In this model, γ-TuRCs are localised to the cis-Golgi and microtubules are nucleated preferentially towards the axon, generating a polarised microtubule network within the soma. Plus-end-associated Kinesin-2 guides the growing plus ends of the nucleated microtubules along and towards the plus ends of the polarised microtubule network, and thus guides them towards the axon and away from dendrites (Figure 8A,C). Microtubules that happen to grow towards a dendrite are excluded from entering when the plus-end-associated Kinesin-2 engages with the shaft of a dendritic microtubule of opposite polarity and thus exerts a backward stalling force on the growing plus end (Figure 8B,C). Importantly, this model explains how minus-end-out microtubule polarity is maintained within proximal dendrites.

**Figure 8.**
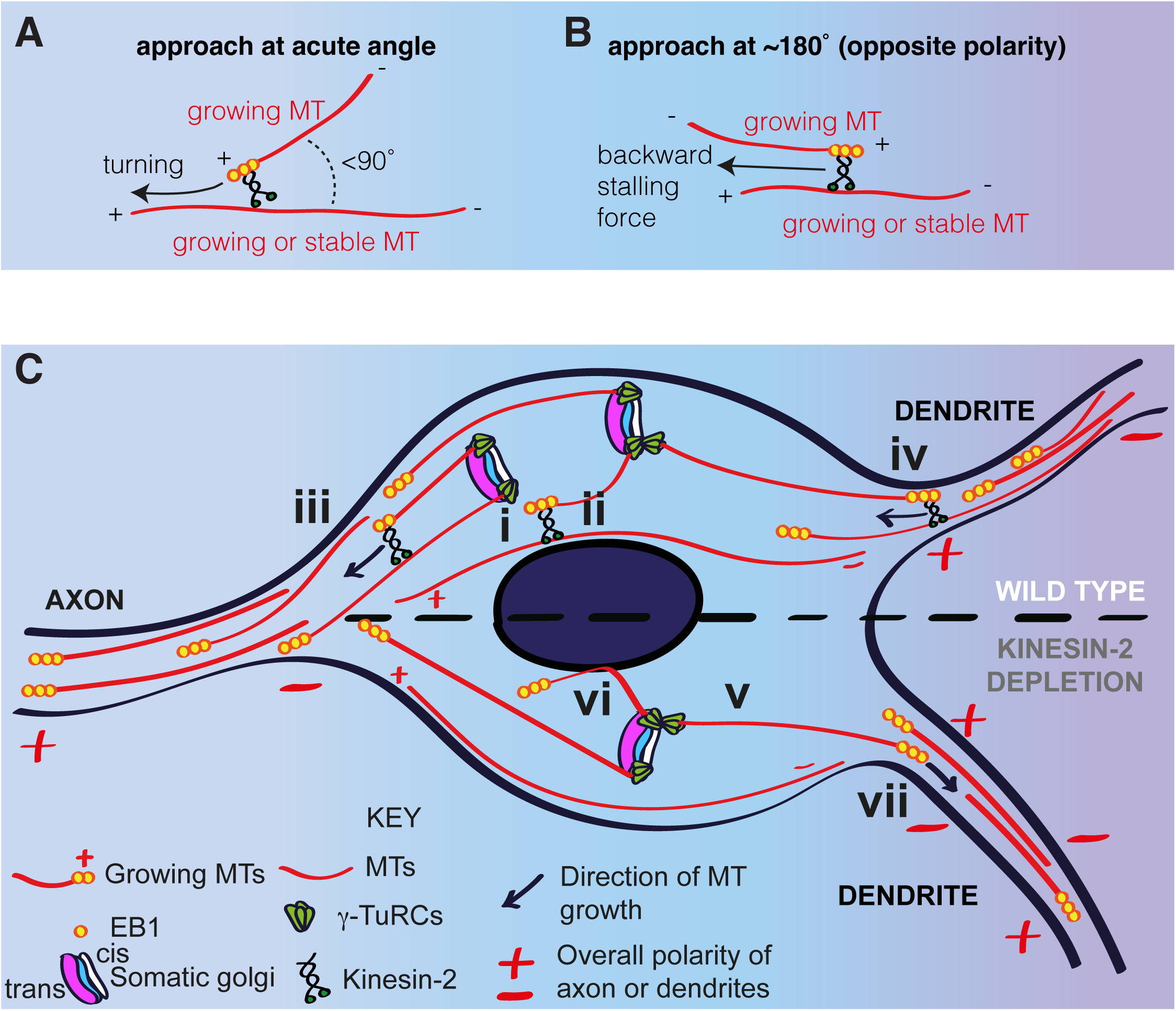
A model for microtubule nucleation and organisation within the soma of da neurons. (**A,B**) Diagrams showing how plus-end-associated Kinesin-2 is predicted to induce turning of a growing microtubule along another microtubule when it approaches the other microtubule at an acute angle (A), or plus-end stalling when the growing microtubule approaches the other microtubules at an angle of ∼180° (B). (**C**) A model depicting microtubule nucleation and growth within the soma of wild type (upper section) or Kinesin-2 depleted (lower section) da neurons. γ-TuRCs localise to the cis-Golgi (white) and nucleate microtubules preferentially towards the axon (i), creating a polarised microtubule network. In wild type neurons, plus-end-associated Kinesin-2 guides growing microtubules along this network towards the axon (ii). Growing microtubules readily enter axons (iii) but are excluded from entering dendrites (iv), because plus-end-associated Kinesin-2 engages with microtubules of opposite polarity, exerting a backward stalling force on the growing microtubule. In Kinesin-2 depleted neurons, microtubules nucleated from the somatic Golgi grow in straight lines (v) until contact with the nuclear envelope (vi) or the cell cortex (not depicted). Microtubules that approach dendrites can now readily enter (vii) resulting in a loss of overall minus-end-out microtubule polarity.

## Discussion

Understanding of how neuronal microtubules are generated and organised is a matter of much interest and importance. In this paper, we have used *Drosophila* da neurons as an *in vivo* model system to address these questions. Our results have led us to propose a model that describes how microtubules are generated and organised within the soma, which also impacts the microtubule populations within proximal axons and dendrites. This model comprises multiple elements. Firstly, the model states that microtubules are nucleated asymmetrically from the somatic Golgi, growing initially with a preference towards the axon and away from dendrites. Secondly, plus-end-associated Kinesin-2 helps guide these growing microtubules towards the axon and away from dendrites along a polarised network of microtubules within the soma. Thirdly, plus-end-associated Kinesin-2 also helps prevent any microtubules that do approach the entrance to a dendrite from entering. We propose that a combination of these mechanisms is essential to maintain minus-end-out polarity in proximal dendrites, a key feature that distinguishes dendrites from axons.

We have shown that endogenously tagged γ-tubulin23C-GFP, which is a proxy for the major catalysts of microtubule nucleation, γ-TuRCs, localises predominantly to the Golgi stacks within the soma of da neurons, and to a few Golgi outposts within the proximal branches of class IV da neurons. Our data shows that the ability of γ-tubulin to localise to these Golgi structures does not depend upon either Cnn or Plp, both of which have been implicated in recruiting γ-TuRCs to Golgi outposts within dendrites of class I and class IV da neurons, respectively (Ori-McKenney et al., 2012; Yalgin et al., 2015). Thus, γ-tubulin, presumably in the form of γ-TuRCs, must be recruited to the Golgi by another tethering protein, perhaps by one of the large coiled-coil “Golgin” proteins that project from specific parts of the Golgi (Munro, 2011).

After showing that γ-tubulin-GFP localised to the somatic Golgi, we tracked EB1-GFP comets from the Golgi and found that they grew with an initial directional preference towards the axon, suggesting of asymmetric nucleation. This analysis was more robust under normal conditions than after cooling-warming cycles, because more comets could be analysed (it was difficult to keep the imaging plane consistent during temperature shifting, due to temperature-induced glass expansion/contraction). Thus, while an equal number of Golgi-derived comets (4) grew towards and away from the axon after warming in Movie S3, we believe this perceived lack of asymmetry is most likely due to chance, especially as a fraction of comets can grow in more random directions even under normal conditions (Figure 6C). We therefore propose there is a genuine preference for growth of Golgi-derived microtubules towards the axon. This may be explained, at least in part, by the asymmetric localisation of γ-tubulin at the somatic Golgi stacks. Using markers of cis- or trans-Golgi, we have found that γ- tubulin localises preferentially to the cis-Golgi. This is also true in mammalian cells, where the somatic Golgi has been shown to act as an asymmetric MTOC in various cell types, such as migrating fibroblasts and different types of cultured mammalian cells in interphase (Rios, 2014). It has been proposed that asymmetric microtubule nucleation is due to the presence of CLASP proteins at the trans-Golgi, which have been suggested to bind and stabilise the plus ends of microtubules that are nucleated by γ-TuRCs at the cis-Golgi (Vinogradova et al., 2009). Whether this occurs in *Drosophila* neurons remains to be explored. *Drosophila* CLASP is known to be a plus-tip protein within axons (Beaven et al., 2015) and is required for axon guidance (Lee et al., 2004), but to our knowledge a role for CLASP at the Golgi in any *Drosophila* cell type has not yet been reported. In mammalian cells, directional microtubule nucleation is thought to be important for cell motility in fibroblasts (Efimov et al., 2007) and for polarised pseudopod formation during invasion of cancer cells (Wu et al., 2016). Our data suggest that asymmetric nucleation at the somatic Golgi in *Drosophila* da neurons may establish a polarised microtubule network within the soma and is important for ensuring that plus-end-out microtubules enter axons rather than dendrites.

It is possible that a fraction of comets that originate from the somatic Golgi represent microtubules that were re-growing after partial depolymerisation, where the re-growth event happened to overlap a Golgi stack. However, several pieces of evidence suggest that the majority of the Golgi-derived comets represent nucleation events. First, γ-tubulin accumulates strongly at the somatic Golgi and a high accumulation of γ-TuRCs at a particular site is normally indicative of a nucleation site, such as when γ-TuRCs accumulate at mitotic centrosomes. Second, we have found that comets emerge repeatedly from the Golgi in a similar direction, strongly suggestive of repeated nucleation events. Third, we believe it is unlikely that all of the comets emerging from the Golgi after a cooling-warming cycle represent microtubules that had not fully depolymerised and that just happened to re-grow close to the Golgi. Moreover, comets that emerged from the Golgi a significant amount of time after warming did not emerge in the same direction as those that had grown immediately after warming. This strongly suggests they are not the same microtubule re-growing after depolymerisation but represent new nucleation events. Direct imaging of microtubules could help assess the degree of overall microtubule depolymerisation induced by cooling, although cold-resistant microtubules that may remain after cooling could impede the ability to visualise depolymerisation of dynamic microtubules, especially if the dynamic microtubules had originally grown alongside the stable microtubules. Increasing the cooling time before inducing regrowth may cause the depolymerisation of all microtubules, and this could be tested by using fluorescent markers of microtubules or by fixing and immunostaining the samples. These experiments have limitations, however, as keeping the animal alive for long enough to depolymerise all microtubule may be challenging and fixing and staining experiments do not allow one to observe the same neuron before, during, and after cooling. In summary, when we consider all of our data as a whole, we believe the most parsimonious conclusion is that microtubules are nucleated asymmetrically from the somatic Golgi.

It has previously been shown that centrosomes in *Drosophila* neurons do not act as MTOCs (Nguyen et al., 2011), and our observations that EB1-GFP comets do not emerge radially from a single source support this conclusion. In other cell types the somatic Golgi can “compete” with centrosomes for the ability to organise microtubules, where decreasing nucleation from one organelle increases nucleation at the other (Gavilan et al., 2018; Ríos et al., 2004; Wu et al., 2016). It is therefore possible that the lack of microtubule nucleation at centrosomes in *Drosophila* neurons enables microtubules to be nucleated at the Golgi. In mammalian neurons the centrosome is deactivated after the first few days in culture (Stiess et al., 2010). An intriguing possibility is that this deactivation allows the Golgi to organise microtubules, which may be necessary to ensure correct microtubule polarity during later neuronal development, although nucleation from the Golgi in mammalian neurons has yet to be reported.

Our analysis was carried out in third instar larvae where the neurons are fully developed and functional. Axons have already extended and made connections in the central nervous system and dendritic arbors are largely established. It is thus somewhat surprising that microtubules continue to be nucleated within the soma. It is possible that somatic microtubules are necessary for transport of cargos to axonal and dendritic entry sites. Given that Golgi-derived microtubules grow preferentially towards the axon and that somatic Golgi stacks are distributed throughout the soma, there is likely an overall microtubule polarity within the soma itself, with more plus ends pointing towards the axon. Even small biases in the orientation of microtubules within a network can be important for polarised distribution of components, such as in *Drosophila* oocytes (Zimyanin et al., 2008). A similar bias within neuronal soma could be important for directing molecules into axons and dendrites. Perhaps this asymmetric microtubule network within the soma, as well as the microtubules within the axon, need constant renewal. There is evidence that molecular motors can damage microtubules (Triclin et al., 2018) and so the large amount of motor-driven transport that occurs within neurons may necessitate microtubule renewal from the soma.

A key feature of our model is the action of plus-end-associated Kinesin-2. Our Kap3 RNAi data suggests that Kinesin-2 is required to guide growing microtubules along a polarised microtubule network within the soma towards the axon and away from dendrites, although future experiments with dual-colour imaging of microtubules and EB1-GFP comets will help support this conclusion. Our data also suggests that plus-end-associated Kinesin-2 prevents outward growing plus ends of somatic microtubules from entering dendrites. Within the soma, we envisage that when a somatic microtubule approaches another somatic microtubule at an acute angle i.e. less than 90°, it can readily turn and be guided along that microtubule (Figure 8A,C). These turning events also occur at dendritic branch points (Mattie et al., 2010; Weiner et al., 2016) and have been recapitulated *in vitro* (Chen et al., 2014; Doodhi et al., 2014). One study showed that the *in vitro* turning events also occured at obtuse angles (>90°), where the growing microtubule buckled to allow the plus end to remain in contact with the ‘rail’ microtubule. This could presumably also occur within the soma of da neurons. The frequency of guidance events, however, decreases significantly as the angle increases (Doodhi et al., 2014). In contrast to the microtubule collision events that will occur with variable angles within the neuronal soma, when outward growing somatic microtubules try to grow into a dendrite they will always encounter dendritic microtubules at angles close to 180° (Figure 8B,C), because most dendritic microtubules are oriented minus-end-out and also because of the limited space available at dendrite entry points. Kinesin-2 motors on the plus end of the outward growing somatic microtubules will engage with the shafts of the minus-end-out dendritic microtubules and generate a backwards force when attempting to ‘walk’ towards the plus end of the dendritic microtubule i.e. in a direction opposite to which the plus end of the somatic microtubule is attempting to grow. This backward force would presumably lead to stalling of the plus end and thus depolymerisation of the microtubule. It is also possible that Kinesin-2 motors slide the plus end along the dendritic microtubule back into the soma, but this sliding action requires the growing microtubule to buckle and thus requires sufficient space (Doodhi et al., 2014). Thus, buckling may be restricted at the narrow dendrite entry site, and so presumably depolymerisation occurs instead. Kinesin-2 motors should be sufficient to induce stalling of microtubule growth, because growing microtubules generate a force of ∼3- 4pN (Dogterom and Yurke, 1997), while a single Kinesin-2 motor generates a force of ∼5pN (Schroeder et al., 2012). Depolymerisation upon stalling could be equivalent to how depolymerisation is induced when microtubules grow against immovable objects (Janson et al., 2003). Importantly, this model elegantly explains why microtubules from the soma are prevented from growing into the dendrites while dendritic microtubules can readily grow into the soma. Other mechanisms, such as a high concentration of a microtubule depolymerase at dendritic entry sites could not distinguish between these two events. In contrast, our model predicts that dendritic microtubules can grow readily into the soma because they do not frequently encounter a microtubule with opposite polarity, except for the occasional somatic microtubule attempting to grow into a dendrite.

Our model explains how minus-end-out microtubule polarity is maintained within proximal dendrites, but what is the origin of minus-end-out microtubules? These microtubules could have been pushed out from the soma minus end first by motor-based sliding, a mechanism known to promote initial axon outgrowth in cultured *Drosophila* neurons (Lu et al., 2013, 2015). Alternatively, or in conjunction with this, they could have been nucleated at sites within dendrites and then have grown back towards the soma. Retrograde microtubule growth events do occur within the dendrites of da neurons, and some of these growth events originate from Golgi outposts (Ori-McKenney et al., 2012; Yalgin et al., 2015; Zhou et al., 2014) and from early endosomes at branchpoints (Weiner et al., 2020). It was shown that γ-tubulin co-localises with early endosome markers at class I branchpoints and that this localisation relies on Wnt signalling proteins (Weiner et al., 2020). Our observation that endogenously-tagged γ-tubulin localises to branchpoints supports this conclusion. Moreover, it was recently shown that an endosome-based MTOC tracks the growing dendritic growth cone within *C. elegans* PVD neurons and nucleates microtubules that travel back towards the soma (Liang et al., 2019). The authors also suggest that a similar process may occur in newly developing *Drosophila* class I neurons. We also observe γ-tubulin within dendritic bubbles and dendritic stretches, but whether γ- tubulin is also recruited to these regions by endosomes, or by cytosolic accumulations of γ-TuRC tethering proteins such as Cnn or Plp, will need to be addressed in future studies. While our data suggest that γ-TuRCs are absent from the majority of Golgi outposts, they appear to be present at a few proximal Golgi outposts within class IV neurons. Moreover, microtubule growth events from Golgi outposts could be independent of γ-TuRCs; other non-γ-TuRC microtubule binding proteins can promote microtubule nucleation *in vitro* (Roostalu and Surrey, 2017), and ectopic Golgi outposts within the axons of nudE mutants promote microtubule growth but not via γ- TuRCs (Arthur et al., 2015; Yang and Wildonger, 2019). Thus, there are several potential mechanisms to generate minus-end-out microtubules within dendrites, that will need to be explored further in future.

It will be interesting in future to examine whether the plus ends of somatic microtubules can grow into dendrites more readily during early neuronal development, when there may be fewer minus-end-out microtubules within dendrites. If true, there must come a developmental point at which the balance shifts and microtubules are prevented from growing into dendrites. This shift could occur in several different ways, including upregulating the number of retrograde microtubules nucleated within dendrites, upregulating Kinesin-2, or changing the direction of microtubule nucleation within the soma. These are all interesting possibilities that are now open for investigation in the future.

## Acknowledgements

This work was supported by a Wellcome Trust and Royal Society Sir Henry Dale Fellowship (105653/Z/14/Z) and an Isaac Newton Trust Research grant (18.23(p)) awarded to PTC, and an Association pour la Recherche sur le Cancer grant (PJA 20181208148) awarded to AG. We thank other members of the Conduit lab for their invaluable input and critical reading of the manuscript. We thank Matthias Landgraf, Melissa Rolls, Adrian Moore, Jill Willdonger for fruitful discussion during the course of the project. We thank Sean Munro and Alison Gillingham for advice on the Golgi and providing us with the Arl1 antibody. We thank Darren Williams and Bing Ye for fly lines that were used in this study. The work benefited from use of the Imaging Facility, Department of Zoology, supported by Matt Wayland and a Sir Isaac Newton Trust Research Grant (18.07ii(c)).

## Author Contributions

PTC and AM designed the study, analysed data and wrote the manuscript. AM performed the vast majority of experiments; PB imaged and quantified ManII-mCherry-labelled Golgi outposts; PTC performed some of the EB1-GFP comet imaging. FB and AG supplied the γ-tubulin-ssss-eGFP fly line.

## Materials and methods

### Contact for Reagent and Resource Sharing

Further information and requests for resources and reagents should be directed to and will be fulfilled by the Lead Contact, Paul Conduit.

### Experimental Model and Subject Details

All fly strains were maintained at 18 or 25°C on Iberian fly food made from dry active yeast, agar, and organic pasta flour, supplemented with nipagin, propionic acid, pen/strep and food colouring.

#### *Drosophila melanogaster* stocks

The following fluorescent alleles were used in this study: γ-tubulin23C-sfGFP (Tovey et al., 2018), γ-tubulin23c-eGFP (this study), sfGFP-Cnn-P1 (this study), sfGFP-Cnn-P2 (this study), UAS-mCD8-mRFP (BL 27392), UAS-EB1-GFP (BL 35512), UAS-ManII-mCherry (Yuh-Nung Jan), and UAS-ManII-GFP (BL 65248). The following Gal4 lines were used in this study: ppk-Gal4 Chr II (B32078) and Chr III (BL 32079) and 221-Gal4 (BL 26259). The following mutant alleles were used in this study: *plp^5^* (BL 9567), *plp^s2172^* (BL 12089), *cnn^f04547^* (Exelixis at HMS), ppkCD4-tdGFP (BL 35842), KAP RNAi (VDRC 45400 GD), γ-tubulin37C RNAi (BL32513), Klp64D RNAi (VDRC 45373) UAS-Dicer 2 (BL 24646).

For examining the endogenous localisation of γ-tubulin23C we used flies expressing γ-tubulin23C-sfGFP and γ-tubulin23C-eGFP i.e. 2 copies of γ-tubulin23C-GFP, either alone or in combination with one copy of UAS-mCD8-RFP expressed under the control of one copy of either 221-Gal4 or ppk-Gal4. For examining the localisation of Cnn, we used flies expressing two copies of either sfGFP-Cnn-P1 or sfGFP-Cnn-P2 either alone or in combination with one copy of UAS-mCD8-RFP expressed under the control of one copy of either 221-Gal4 or ppk-Gal4. For examining the localisation of Plp in relation to medial Golgi, we used flies with one copy of UAS-ManII-GFP expressed under the control of one copy of either 221-Gal4 or ppk-Gal4. For examining the localisation of γ-tubulin23C in the absence of Cnn or Plp, we used flies expressing two copies of γ-tubulin23C-(sf/e)GFP (as above) in a *cnn^f04547^*/*cnn ^f04547^* or *plp^5^*/*plp^s2172^* mutant background. For examining microtubule dynamics in relation to the Golgi we used flies with one copy of UAS-EB1-GFP and one copy of UAS-ManII-mCherry, both expressed under the control of one copy of 221-Gal4. For examining microtubule dynamics in control neurons we used flies with one copy of UAS-EB1-GFP and one copy of UAS-γ-tubulin37C RNAi, with or without one copy of UAS-Dicer 2, expressed under the control of one copy of 221-Gal4. For examining microtubule dynamics in Kap3 RNAi neurons we used flies with one copy of UAS-EB1-GFP, one copy of UAS- Dicer 2 and one copy of UAS-Kap3 RNAi expressed under the control of one copy of 221-Gal4.

### Method Details

#### DNA cloning

5-alpha Competent *E. coli* (High Efficiency) (NEB) cells were used for bacterial transformations, DNA fragments were purified using QIAquick Gel Extraction Kits (Qiagen), plasmid purification was performed using QIAprep Spin Miniprep Kits (Qiagen). Phusion High-Fidelity PCR Master Mix with HF Buffer (ThermoFisher Scientific) was used for PCRs.

#### Generating endogenously-tagged fly lines

All endogenously-tagged lines were made using CRISPR combined with homologous recombination, by combining the presence of a homology-repair vector containing the desired insert with the appropriate guide RNAs and Cas9. The *γ-tubulin23C-eGFP* allele was generated by inDroso by initially inserting an SSSS-eGFP-3’UTR-LoxP- 3xP3-dsRED-Lox P cassette before the selection markers were excised. The multi-serine insert acts as a flexible linker between γ-tubulin23C and eGFP. The following guide RNA sequences were used to cut either side of the 3’UTR: AGTCGATC|TGTGACCAGCGC and TTATGGTT|AATGTCGACTTG. The sfGFP-Cnn-P1 (insertion of sfGFP at the start of exon 1a) and sfGFP-Cnn-P2 (insertion of sfGFP at the start of exon 1b) alleles were generated within the lab following a similar approach to that used previously (Tovey et al., 2018). For GFP-Cnn-P1, flies expressing a single guide RNA (GTCGTGTTTAGACTGGTCCATGGG) were crossed to nos-Cas9 expressing females and the resulting embryos were injected by the Department of Genetics Fly Facility, Cambridge, UK, with a pBluescript plasmid containing sfGFP and linker sequence (4X GlyGlySer) flanked on either side by 1.5kb of DNA homologous to the *cnn* genomic locus surrounding the 5’ end of the appropriate coding region. The homology vectors were made by HiFi assembly (NEB) of PCR fragments generated from genomic DNA prepared from nos-Cas9 flies (using MicroLYSIS, Microzone) and a vector containing the sfGFP tag (DGRC, 1314). For GFP-Cnn-P2 flies, both the guide (GTCGTTAAATGAAACATAGAATA) and the homology vector were injected into nos-Cas9 embryos (BL54591) by Rainbow Transgenic Flies, Inc. Camarillo, CA 93012, USA. F1 and F2 males were screened by PCR using the following primers: for sfGFP-Cnn-P1: forward primer: AAAGTTAACTATTTGAGGACCTCCCATGGTGTCCAAGGGCGAGGAG; reverse primer:

CCCGCAAAACCTGTTTAGACTGGTCGGATCCGCCGCTACCTCCGCTTCCACCGGAACCTCCCTTGTACAGCTCATCCATGCC; for sfGFP-Cnn-P2: forward primer: GCAAATGTTAAATGAAAGATACAATATGGTGTCCAAGGGCGAGGAG; reverse primer:

GTGAGGTAGATCGAAAGATACCCGCGGATCCGCCGCTACCTCCGCTTCCACCGGAACCTCCCTTGTACAGCTCATCCATGCC.

#### Antibodies

The following primary antibodies were used: anti-GFP mouse monoclonal at 1:250 (Roche, 11814460001), anti-Cnn Rabbit polyclonal raised against first 660aa of Cnn-P1 (which includes amino acids 35-632 of Cnn-P2) at 1:1000(Lucas and Raff, 2007), anti-Plp rabbit polyclonal at 1:500 (Martinez-Campos et al., 2004), Alexa-647 conjugated HRP polyclonal at 1:500 (Jackson), anti-Arl1 chicken polyclonal at 1:200 (gift from Sean Munroe (Torres et al., 2014)), and anti-GM130 rabbit polyclonal at 1:300 (Abcam). The following secondary antibodies were used: ATTO-488 GFP- booster at 1:200 (Chromotek), Alexa-488 anti-Mouse at 1:500 (ThermoFisher), Alexa-547 anti-Rabbit at 1:500 (ThermoFisher).

#### Immunostaining

Larvae were dissected as described previously (Broadie and Bate, 1993). The fillet preparations were fixed in freshly prepared 4% formaldehyde for 20 minutes at room temperature and were then washed four times for 10 minutes in PBST (PBS + 0.1% TritonX-100). The perparations were then blocked in PBST + 5% BSA for 1 hour at room temperature and incubated with the appropriate primary antibodies diluted in PBST overnight at 4°C. After washing in PBST for ∼8 hours, changing washes every 30-45 minutes, samples were incubated in secondary antibodies diluted in PBST overnight at 4°C. The fillet perparations were then washed for ∼8 hours, changing washes every 30-45 minutes in PBST before mounting in Moviol. They were stored at −20°C and imaged within a week. Neurons within segments A2 to A6 were imaged; we observed no difference in the staining patterns between these segments.

#### Fixed and live cell imaging

Imaging of all samples except for those expressing EB1-GFP in neurons or GFP-Cnn-P1 in embryos was carried out on an Olympus FV3000 scanning inverted confocal system run by FV-OSR software using a 60X 1.4NA silicon immersion lens (UPLSAPO60xSilcon). For live samples, wandering L3 larvae were placed in a drop of glycerol and flattened between a slide and a 22X22mm coverslip, held in place by double-sided sticky tape, and imaged immediately. Imaging of EB1-GFP within da neurons and GFP-Cnn-P1 within embryos was performed on a Leica DM IL LED inverted microscope controlled by μManager software and coupled to a RetigaR1 monochrome camera (QImaging) and a CoolLED pE-300 Ultra light source using a 63X 1.3NA oil objective (Leica 11506384). For EB1-GFP imaging, larvae were dissected in live imaging solution (ThermoFisher) to remove the majority of their tissues and mounted in Schneider’s medium supplemented with FBS and pen/strep between a slide and 22X22mm coverslip held in place with tape and imaged immediately. When using the CherryTemp, the larvae were held between the CherryTemp chip and a 22X22mm coverslip. The CherryTemp software was used to rapidly change the temperature of the liquid within the flow chamber from 20°C to 5°C after an initial 45s of imaging. The temperature was then rapidly changed back to 20°C after a further 180s and the neurons were imaged for a further 135s. For EB1-GFP imaging, single Z-plane images were acquired every 5 seconds; for EB1-GFP/ManII- mCherry dual imaging single Z-plane images were acquired every 3 seconds. All images were processed using Fiji (ImageJ). EB1 comets were tracked using the Manual Tracking plugin in Fiji.

#### Quantification and Statistical Analysis

Statistical analysis and graph production were performed using GraphPad Prism. To determine punctate versus diffuse localisation of γ-tubulin-GFP at branchpoints of class I da neurons, we visually categorised each branchpoint into one or other group depending on whether the puncta were clear and obvious or not. If they were clear and obvious, then they were categorised as puncta. If puncta and diffuse patches did co-exist we defined them as puncta. To quantify the number of Golgi outposts and γ- tubulin-GFP puncta per 100μm dendrite we measured the total length of dendrites in ImageJ using the segmented line tool and marked and counted the number of Golgi outposts and γ-tubulin-GFP puncta. We then calculated their number per 100μm dendrite. We did this across multiple images of different neurons. We analysed 14 class I neurons with a total dendritic length of 8525μm and a total number of 183 branchpoints. We analysed 10 proximal and 7 distal images of class IV da neurons with a total dendritic length of 3619μm and 4489μm, respectively, and a total number of 47 and 179 branchpoints, respectively.

For EB1-comet analysis, we excluded comets that were present within the first timepoint (and thus may not represent newly growing microtubules). The angle of initial comet growth was measured using the angle tool within ImageJ, by drawing lines from the axon entry site to the comet origin and then along the initial linear path of the comet. For the frequency distribution of comet angles in Figure 6B, the negative angles were made positive, so as not to distinguish between comets growing either side of the axon entry site, creating a distribution between 0° and 180°. A total of 163 comets from 59 Golgi stacks from 7 cells were analysed.

For the vector analysis in Figure 6C,D we used both positive and negative angles (between −180° and 180°). Initially, a vector for each comet was generated with a length of 1 by calculating cosine(angle) (Y value) and sin(angle) (X value). Taking in turn each Golgi stack that generated more than one comet, the vectors of each comet were added together to generate a resultant vector. These resultant vectors were then normalised by dividing their X and Y values by the number of comets originating from the Golgi stack, such that the maximum length of the resultant vector would be 1, irrespective of comet number. These normalised resultant vectors were plotted as circles on a scatter plot, with the size of each circle reflecting the number of comets originating from that particular Golgi stack. 44 Golgi stacks were analysed from 7 neurons. We generated random datasets by replacing each comet angle with a randomly generated number between −180 and 180 to show what results could be expected if the angles of comets originating from Golgi stacks were independent of each other and of the position of the axon (thus each random dataset was generated using 44 hypothetical Golgi stacks). We generated three random datasets, the resultant vector length distributions of which are included in Figure 6D.

One-way Chi-squared tests were used to determine whether frequency distributions were significantly different from the distribution expected by chance i.e. if comet angles were random; a binomial probability calculator was used to determine the chance of observing the skewed distribution between upper (towards the axon) and lower (away from the axon) quadrants in the resultant vector scatter plot in Figure 6C.

To assess whether comets that originated within the soma approached and entered axons or dendrites, we defined a region (∼0.5μm wide) across the entrance to the axon or dendrite; any comets that entered this region were scored as comets that had approached; any comets that crossed this region were scored as comets that entered. For control neurons, we analysed a total of 666 comets (across 13 movies) that had originated during the movies within the soma (average of 10.0 comets per minute); for Kap3 RNAi we analysed a total of 1058 comets (across 9 movies) (average of 20.0 comets per minute). Due to the labour-intensive nature of manual tracking we did not track all comets within the soma of Klp64D RNAi neurons; we instead focussed only on comets that approached dendrites (a total of 80 comets from 10 movies). To assess the polarity of dynamic microtubules within the proximal dendritic regions, we included both comets that originated within the proximal dendrite and those that grew into the dendrite from the soma. For control neurons, we analysed a total of 252 comets from 13 cells; for Kap3 RNAi we analysed a total of 338 comets from 9 cells. The overall proportion of comets that either turned, approached, entered or had anterograde polarity was plotted in Figure 7C-F along with the 95% confidence interval for each dataset. Datasets were compared using one-way Chi-squared tests.

## Supplementary Figure legends

**Figure S1 – related to Figure 1.**
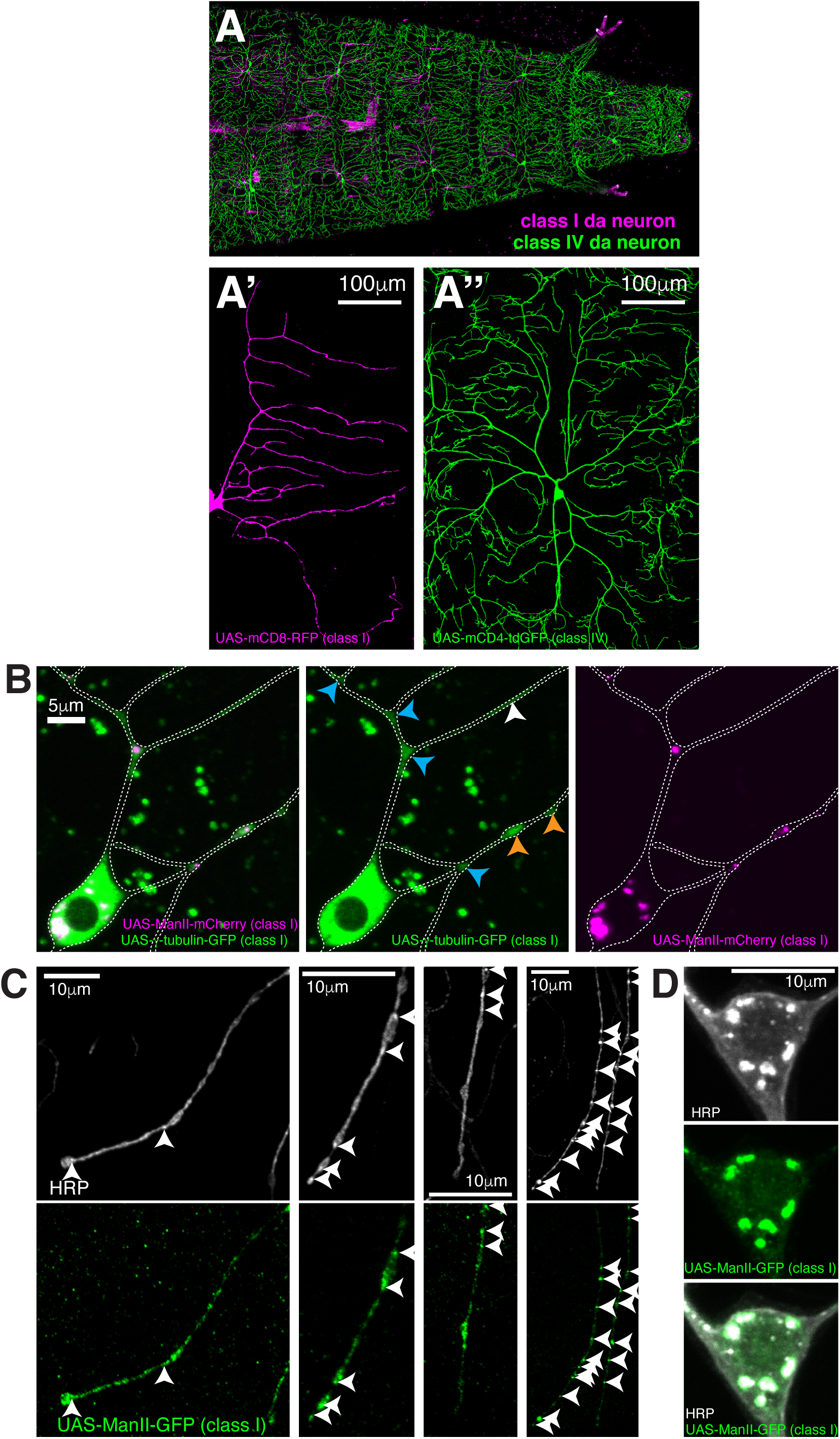
Ectopically expressed γ-tubulin-GFP is strongly enriched in branchpoints and dendritic bubbles and ectopically expressed ManII is not a reliable marker of Golgi outposts in class I da neurons. (**A**) A fluorescent confocal image of the anterior region of a living 3^rd^ instar larva where mCD8-RFP and mCD4-tdGFP are expressed in class I (magenta) and class IV (green) neurons, respectively. (**A’,A’’**) Enlarged images of a class I (A’) and a class IV (A’’) da neuron. (**B**) Fluorescent confocal images of the proximal region of a class I da neuron expressing 221- gal4>UAS-γ-tubulin-GFP (green) and 221-gal4>UAS-ManII-mCherry (magenta) within a living 3^rd^ instar larva. Left panel shows an overlay of the GFP and mCherry channels, middle and right panels show the GFP and mCherry channels, respectively. Blue and orange arrowheads indicate enrichments of ectopically expressed γ-tubulin-GFP within branchpoints and dendritic bubbles, respectively. (**C,D**) Fluorescent confocal images of distal dendrites (C) or a soma (D) from different class I da neurons expressing 221-gal4>UAS-ManII-GFP fixed and immunostained with antibodies against HRP (greyscale) and GFP (green). Arrowheads in (C) indicate HRP puncta that represent internal membrane, at least some of which could be Golgi outposts. Note that UAS-ManII-GFP spreads into regions of the class I dendrites, including dendritic bubbles, that do not contain HRP puncta, suggesting that over-expressing ManII in class I neurons leads to ‘leaking’ of the protein outside of Golgi outposts within dendrites. In contrast, the UAS-ManII-GFP signal always colocalises with the HRP signal within soma (D).

**Figure S2 – related to Figure 3.**
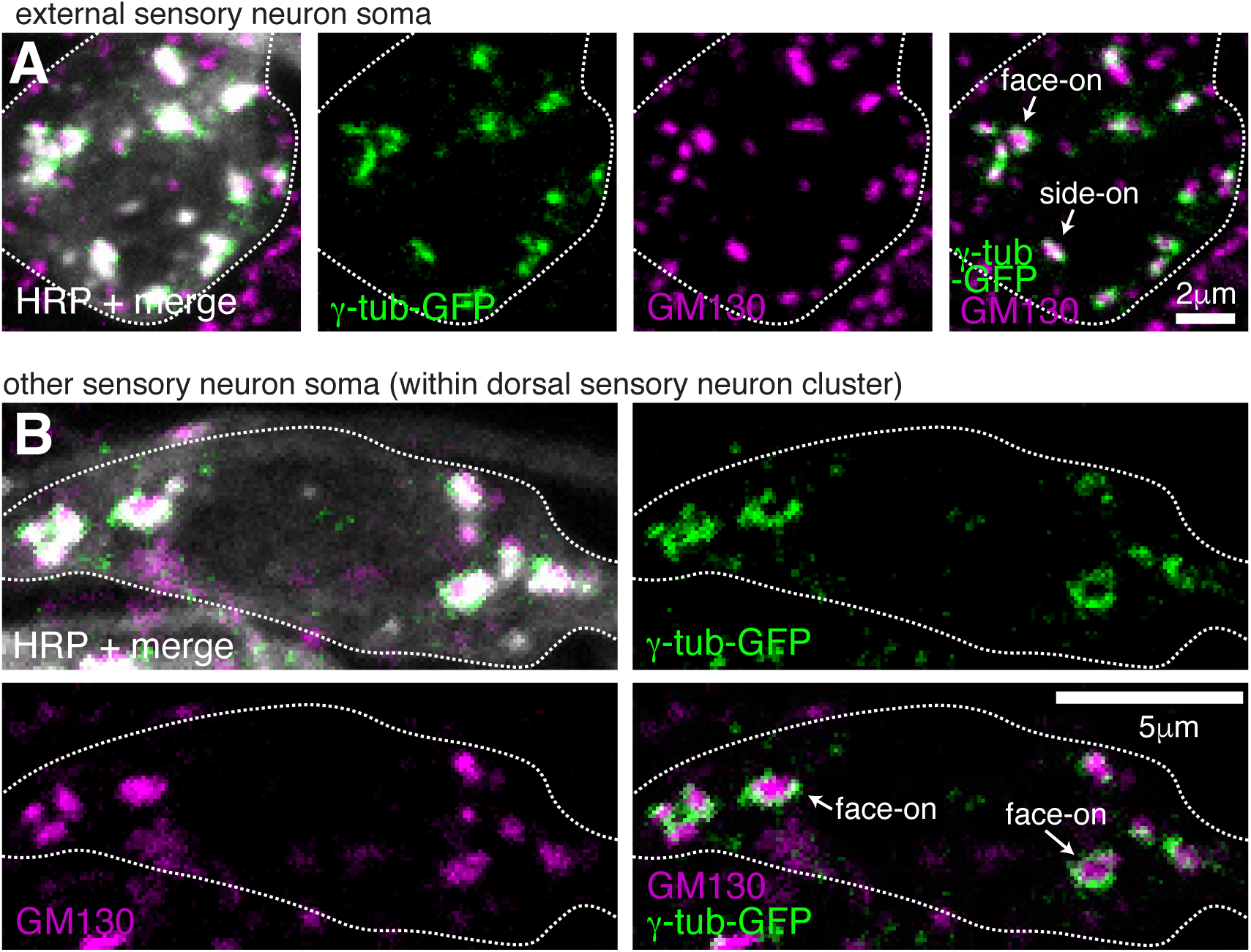
Endogenously-tagged γ-tubulin-GFP localises to the somatic Golgi of sensory neurons. (**A,B**) Confocal images show the soma of an es neuron (A) or another sensory neuron within the dorsal cluster (B) from 3^rd^ instar larva expressing two copies of endogenous γ-tubulin-GFP fixed and immunostained with antibodies against GFP (green), GM130 (magenta) and HRP (greyscale). γ-tubulin-GFP and GM130 signals associate at presumptive side-on stacks (extended GM130 signal) and face-on stacks (round GM130 signal) in a way that suggests γ-TuRCs are recruited to the rims of the cis-Golgi.

**Figure S3 – related to Figure 4.**
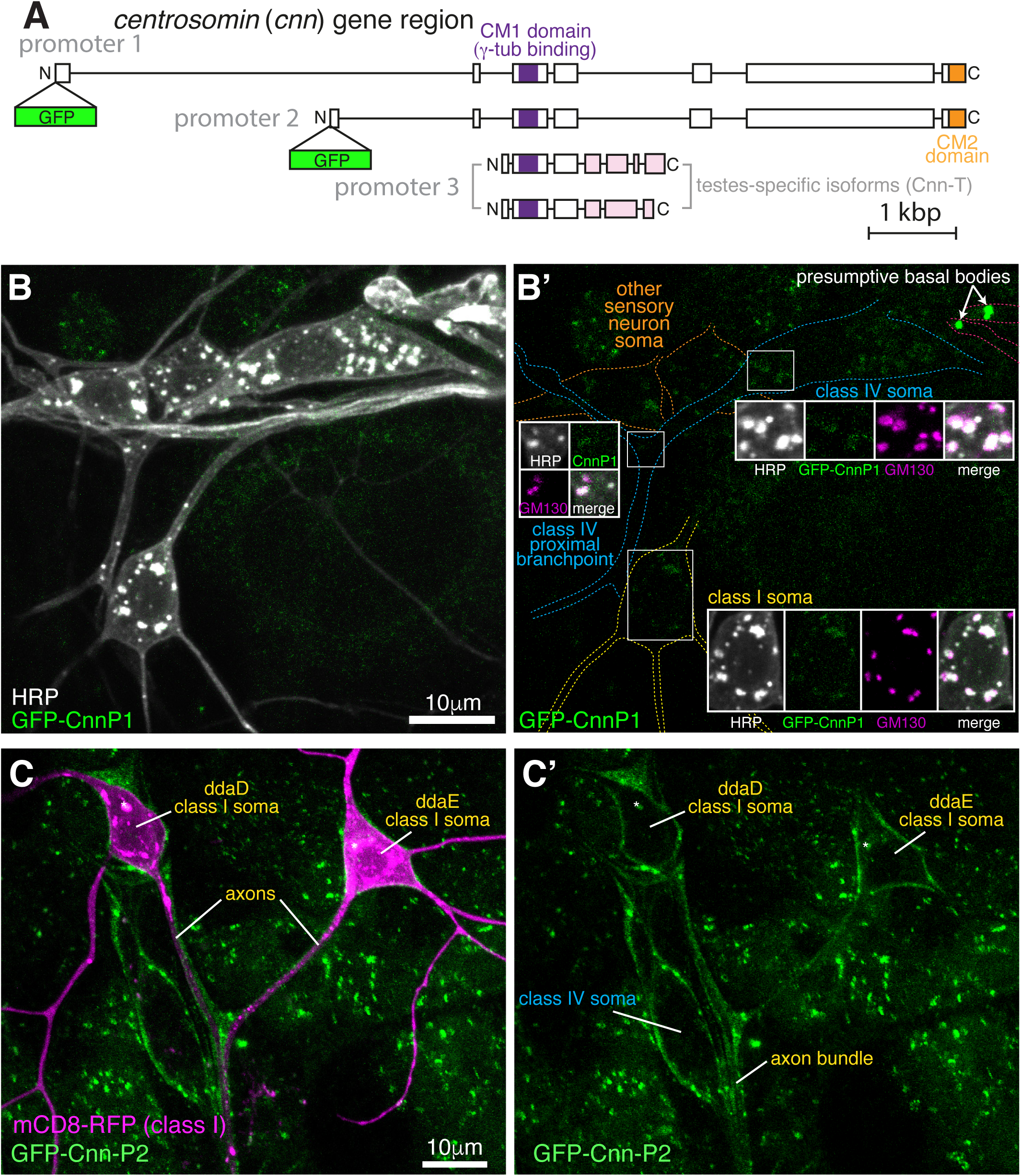
nn-P1 is not strongly associated with the somatic Golgi or Golgi outposts of sensory neurons, while Cnn-P2 is expressed in ensheathing glia. (**A**) A gene diagram of *cnn* to indicate the different Cnn isoforms. Boxed regions are exons, lines are introns. CRISPR combined with homologous recombination was used to insert GFP just after the start codon of either the promoter 1 (P1) or promoter 2 (P2) isoform. Promotor three isoforms are expressed only in testes and so were not analysed in this study. (**B**) Confocal images show the somas and some proximal dendrites of sensory neurons within the dorsal cluster from a 3^rd^ instar larva expressing endogenously-tagged GFP-Cnn-P1 and immunostained with nanobodies against GFP covalently coupled to ATTO-488 (green) and antibodies against HRP (greyscale) and GM130 (magenta). Left panel (B) shows an overlay of the GFP and anti-HRP signals, while the right panel (B’) shows the GFP signal with coloured outlines of the different neurons drawn for clarity, with insets showing all channels, including an overlay, for the class I neuron soma, regions of the class IV neuron soma and a class IV proximal branchpoint that contains Golgi outposts. The GFP-Cnn-P1 signal appears very weakly, if at all, at the somatic Golgi and Golgi outposts marked by HRP and GM130 staining. (**C**) Confocal images show class I da neurons from a living 3^rd^ instar larva expressing 221-Gal4>UAS-mCD8-RFP (magenta) and two copies of endogenously tagged GFP-Cnn-P2. Left panel (C) shows an overlay of the GFP and RFP channels, while the right panel (C’) shows just the GFP channel. Note that the GFP-Cnn-P2 surrounds the axons and somas of the neurons, suggestive of localisation within ensheathing glia. The * star indicates a region of very bright RFP signal that has bled through into the GFP channel.

**Figure S4 – related to Figure 5.**
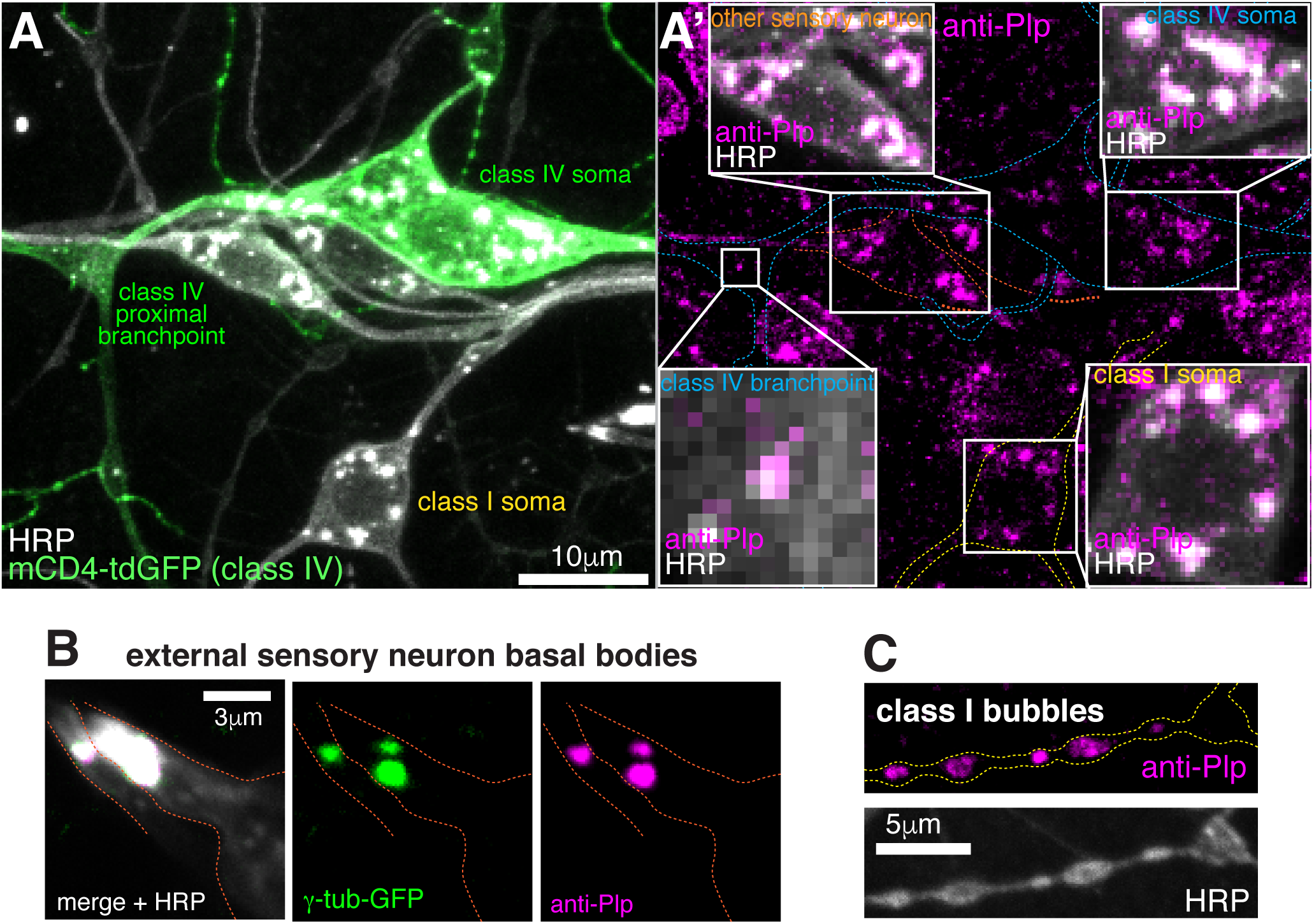
Plp localises to the somatic Golgi, Golgi outposts, basal bodies and class I - specific dendritic bubbles. (**A**) Confocal images show the somas and some proximal dendrites of sensory neurons within the dorsal cluster from a 3^rd^ instar larva expressing ppk-mCD4-tdGFP and immunostained with antibodies against GFP (green), HRP (greyscale) and Plp (magenta). Left panel (A) shows the GFP and anti-HRP channels, while the right panel (A’) shows the anti-Plp channel; insets show an overlay of the anti-Plp and anti-HRP channels. Plp signal colocalises with the HRP staining that marks the multiple well-distributed Golgi stacks within the neuronal soma of each neuron. Plp also localises to some HRP puncta (presumably Golgi outposts) within the proximal branchpoints of class IV neurons. (**B**) Confocal images show parts of the es neurons containing the basal bodies from a 3^rd^ instar larva expressing two copies of endogenously tagged γ- tubulin-GFP and immunostained with antibodies against GFP (green), HRP (greyscale) and Plp (magenta). (**C**) Confocal images show dendritic bubbles of a class I da neuron from a 3^rd^ instar larva immunostained with antibodies against HRP (greyscale) and Plp (magenta).

## Supplementary Movie Legends

**Movie S1 – related to Figure 4.**

**Endogenously-tagged GFP-Cnn-P1 localises as expected to centrosomes within *Drosophila* syncytial embryos.** The Movie was taken using multi-Z-stack time-lapse epifluorescence microscopy and shows Z-projected images through time of a syncytial embryo expressing endogenously-tagged GFP-Cnn-P1. The cycle starts in M-phase and the centrosomes separate when the cycle transitions to the following S-phase. GFP-Cnn-P1 localises to centrosomes during both M-phase, where the signal is rounded, and S-phase, where the signal displays typical Cnn “flares”. Images were collected every 20s and the Movie plays at 20 frames/second.

**Movie S2 – related to Figure 6.**

**EB1-comets emerge from somatic Golgi stacks and initially grow preferentially towards the axon.** The Movie was taken using single-Z time-lapse epifluorescence microscopy and shows a class I da neuron expressing 221-Gal4>UAS-EB1-GFP and 221-Gal4>UAS-ManII-mCherry. The left panel shows an overlay of the GFP and mCherry channels, while the right panel shows the same overlay with the ImageJ- generated EB1-GFP comet tracks drawn onto the images. The axon (A) and dendrites (D) are labelled in the right panel. The tracks are multi-coloured, as is default for ImageJ, and the colours have no reference to comet type. Comets can be seen emerging from Golgi stacks and this can occur repeatedly from the same stack (see bottom right stack). Images were collected every 3s and the Movie plays at 10 frames/second.

**Movie S3 – related to Figure 6.**

**The somatic Golgi stacks nucleate microtubules.** The Movie was taken using single-Z time-lapse epifluorescence microscopy and shows a class I da neuron expressing 221-Gal4>UAS-EB1-GFP and 221-Gal4>UAS-ManII-mCherry. The sample was subjected to a warming-cooling-warming (20°C-5°C-20°C) cycle, as indicated by the red (20°C) and blue (5°C) borders. The left panel shows an overlay of the GFP and mCherry channels, while the right panel shows the same overlay with the ImageJ-generated EB1-GFP comet tracks drawn onto the images. The axon (A) and dendrites (D) are labelled in the right panel. The tracks have been manually post-coloured such that green represent comets emerging from Golgi stacks and purple represent comets emerging from elsewhere within the soma. Comets can be seen emerging from Golgi stacks before cooling, and then all comets stop during the 3 minute cooling period. After warming, comets can be seen emerging from the Golgi stacks, including comets that emerge >1min after warming. Images were collected every 3s and the Movie plays at 10 frames/second.

**Movie S4 – related to Figure 7.**

**Microtubules turn towards the axon and are excluded from entering dendrites.** The Movie was taken using single-Z time-lapse epifluorescence microscopy and shows a control class I da neuron expressing 221-Gal4>UAS-EB1-GFP and 221- Gal4>UAS-γ-tubulin-37c-RNAi. The left panel shows the GFP channel, while the right panel shows the GFP channel with the ImageJ-generated EB1-GFP comet tracks drawn onto the images. The axon (A) and dendrites (D) are labelled in the right panel. The tracks are multi-coloured, as is default for ImageJ, and the colours have no reference to comet type. Comets display turning events, indicating that the growing plus ends of microtubules can be guided along pre-existing microtubules. Comets that approach the axon frequently enter and grow down the axon; In contrast, comets that approach a dendrite entry site do not enter (quantified in Figure 7). Images were collected every 5s and the Movie plays at 10 frames/second.

**Movie S5 – related to Figure 7.**

**Kinesin-2 is required for growing microtubules to turn within the soma and to be excluded from entering dendrites.** The Movie was taken using single-Z time-lapse epifluorescence microscopy and shows a Kinesin-2 RNAi class I da neuron expressing 221-Gal4>UAS-EB1-GFP and 221-Gal4>UAS-Kap3-RNAi. The left panel shows the GFP channel, while the right panel shows the GFP channel with the ImageJ-generated EB1-GFP comet tracks drawn onto the images. The axon (A) and dendrites (D) are labelled in the right panel. The tracks are multi-coloured, as is default for ImageJ, and the colours have no reference to comet type. Comets can be seen emerging from multiple locations within the soma but unlike in control neurons they do not turn unless they encounter the nuclear envelope or cell cortex, suggesting that Kinesin-2 is required to guide growing microtubules towards the axon. Comets that approach the axon still frequently enter and grow down the axon, and, in contrast to control neurons, comets that approach dendrites also readily enter (quantified in Figure 7). This shows that Kinesin-2 is also required for excluding growing microtubules from entering dendrites. Images were collected every 5s and the Movie plays at 10 frames/second.

**Movie S6 – related to Figure 7.**

**Knockdown of Klp64D supports the conclusion that Kinesin2 is required for growing microtubules to be excluded from entering dendrites.** The Movie was taken using single-Z time-lapse epifluorescence microscopy and shows a Kinesin-2 RNAi class I da neuron expressing 221-Gal4>UAS-EB1-GFP and 221-Gal4>UAS- Klp64D RNAi. Images were collected every 5s and the Movie plays at 10 frames/second.

## References

Arthur, A.L., Yang, S.Z., Abellaneda, A.M., and Wildonger, J. (2015). Dendrite arborization requires the dynein cofactor NudE. J Cell Sci 128, 2191–2201.

Baas, P.W., Karabay, A., and Qiang, L. (2005). Microtubules cut and run. Trends Cell Biol 15, 518–524.

Bahi-Buisson, N., Poirier, K., Fourniol, F., Saillour, Y., Valence, S., Lebrun, N., Hully, M., Bianco, C.F., Boddaert, N., Elie, C., et al. (2014). The wide spectrum of tubulinopathies: what are the key features for the diagnosis? Brain 137, 1676–1700.

Beaven, R., Dzhindzhev, N.S., Qu, Y., Hahn, I., Dajas-Bailador, F., Ohkura, H., and Prokop, A. (2015). Drosophila CLIP-190 and mammalian CLIP-170 display reduced microtubule plus end association in the nervous system. Mol Biol Cell 26, 1491–1508.

Bosc, C., Andrieux, A., and Job, D. (2003). STOP Proteins †. Biochemistry-Us 42, 12125–12132.

Broadie, K., and Bate, M. (1993). Activity-dependent development of the neuromuscular synapse during drosophila embryogenesis. Neuron 11, 607–619.

Bucciarelli, E., Pellacani, C., Naim, V., Palena, A., Gatti, M., and Somma, M.P. (2009). Drosophila Dgt6 Interacts with Ndc80, Msps/XMAP215, and γ-Tubulin to Promote Kinetochore-Driven MT Formation. Curr Biol 19, 1839–1845.

Castillo, U. del, Lu, W., Winding, M., Lakonishok, M., and Gelfand, V.I. (2014). Pavarotti/MKLP1 Regulates Microtubule Sliding and Neurite Outgrowth in Drosophila Neurons. Curr Biol 25, 1–6.

Castillo, U. del, Winding, M., Lu, W., and Gelfand, V.I. (2015). Interplay between kinesin-1 and cortical dynein during axonal outgrowth and microtubule organization in Drosophila neurons. Elife 4, e10140.

Chen, J.V., Buchwalter, R.A., Kao, L.-R., and Megraw, T.L. (2017). A Splice Variant of Centrosomin Converts Mitochondria to Microtubule-Organizing Centers. Curr Biol 27, 1928–1940.e6.

Chen, Y., Rolls, M.M., and Hancock, W.O. (2014). An EB1-Kinesin Complex Is Sufficient to Steer Microtubule Growth In Vitro. Curr Biol 24, 316–321.

Cunha-Ferreira, I., Chazeau, A., Buijs, R.R., Stucchi, R., Will, L., Pan, X., Adolfs, Y., Meer, C. van der, Wolthuis, J.C., Kahn, O.I., et al. (2018). The HAUS Complex Is a Key Regulator of Non-centrosomal Microtubule Organization during Neuronal Development. Cell Reports 24, 791–800.

Delphin, C., Bouvier, D., Seggio, M., Couriol, E., Saoudi, Y., Denarier, E., Bosc, C., Valiron, O., Bisbal, M., Arnal, I., et al. (2012). MAP6-F Is a Temperature Sensor That Directly Binds to and Protects Microtubules from Cold-induced Depolymerization. J Biol Chem 287, 35127–35138.

Dogterom, M., and Yurke, B. (1997). Measurement of the Force-Velocity Relation for Growing Microtubules. Science 278, 856–860.

Doodhi, H., Katrukha, E.A., Kapitein, L.C., and Akhmanova, A. (2014). Mechanical and Geometrical Constraints Control Kinesin-Based Microtubule Guidance. Curr Biol 24, 322–328.

Efimov, A., Kharitonov, A., Efimova, N., Loncarek, J., Miller, P.M., Andreyeva, N., Gleeson, P., Galjart, N., Maia, A.R.R., McLeod, I.X., et al. (2007). Asymmetric CLASP- dependent nucleation of noncentrosomal microtubules at the trans-Golgi network. Dev Cell 12, 917–930.

Eisman, R.C., Phelps, M.A.S., and Kaufman, T.C. (2009). Centrosomin: a complex mix of long and short isoforms is required for centrosome function during early development in Drosophila melanogaster. Genetics 182, 979–997.

Farache, D., Emorine, L., Haren, L., and Merdes, A. (2018). Assembly and regulation of γ-tubulin complexes. Open Biol 8, 170266.

Gavilan, M.P., Gandolfo, P., Balestra, F.R., Arias, F., Bornens, M., and Rios, R.M. (2018). The dual role of the centrosome in organizing the microtubule network in interphase. Embo Rep 19, e45942.

Goodson, H.V., and Jonasson, E.M. (2018). Microtubules and Microtubule-Associated Proteins. Csh Perspect Biol 10, a022608.

Grueber, W.B., Jan, L.Y., and Jan, Y.-N. (2002). Tiling of the Drosophila epidermis by multidendritic sensory neurons. Development (Cambridge, England) 129, 2867–2878.

Han, C., Jan, L.Y., and Jan, Y.-N. (2011). Enhancer-driven membrane markers for analysis of nonautonomous mechanisms reveal neuron–glia interactions in Drosophila. Proc National Acad Sci 108, 9673–9678.

Harterink, M., Edwards, S.L., Haan, B. de, Yau, K.W., Heuvel, S. van den, Kapitein, L.C., Miller, K.G., and Hoogenraad, C.C. (2018). Local microtubule organization promotes cargo transport in C. elegans dendrites. J Cell Sci 131, jcs.223107.

Hayward, D., Metz, J., Pellacani, C., and Wakefield, J.G. (2014). Synergy between multiple microtubule-generating pathways confers robustness to centrosome-driven mitotic spindle formation. Dev Cell 28, 81–93.

Hill, S.E., Parmar, M., Gheres, K.W., Guignet, M.A., Huang, Y., Jackson, F.R., and Rolls, M.M. (2012). Development of dendrite polarity in Drosophila neurons. Neural Dev 7, 34.

Jan, Y.-N., and Jan, L.Y. (2010). Branching out: mechanisms of dendritic arborization. Nat Rev Neurosci 11, 316–328.

Janson, M.E., Dood, M.E. de, and Dogterom, M. (2003). Dynamic instability of microtubules is regulated by force. J Cell Biology 161, 1029–1034.

Kapitein, L.C., and Hoogenraad, C.C. (2015). Building the Neuronal Microtubule Cytoskeleton. Neuron 87, 492–506.

Kelliher, M.T., Saunders, H.A., and Wildonger, J. (2019). Microtubule control of functional architecture in neurons. Curr Opin Neurobiol 57, 39–45.

Klinman, E., Tokito, M., and Holzbaur, E.L.F. (2017). CDK5-dependent activation of dynein in the axon initial segment regulates polarized cargo transport in neurons. Traffic 18, 808–824.

Kondylis, V., and Rabouille, C. (2009). The Golgi apparatus: Lessons from Drosophila. Febs Lett 583, 3827–3838.

Lee, H., Engel, U., Rusch, J., Scherrer, S., Sheard, K., and Vactor, D.V. (2004). The Microtubule Plus End Tracking Protein Orbit/MAST/CLASP Acts Downstream of the Tyrosine Kinase Abl in Mediating Axon Guidance. Neuron 42, 913–926.

Liang, X., Kokes, M., Fetter, R., Pickett, M.A., Sallee, M.D., Moore, A.W., Feldman, J., and Shen, K. (2019). Growth Cone-Localized Microtubule Organizing Center Establishes Microtubule Orientation in Dendrites. Biorxiv 841759.

Lin, T., Neuner, A., and Schiebel, E. (2014). Targeting of γ-tubulin complexes to microtubule organizing centers: conservation and divergence. Trends Cell Biol 25, 296–307.

Lu, W., Fox, P., Lakonishok, M., Davidson, M.W., and Gelfand, V.I. (2013). Initial neurite outgrowth in Drosophila neurons is driven by kinesin-powered microtubule sliding. Curr Biol 23, 1018–1023.

Lu, W., Lakonishok, M., and Gelfand, V.I. (2015). Kinesin-1-powered microtubule sliding initiates axonal regeneration in Drosophila cultured neurons. Mol Biol Cell 26, 1296–1307.

Lucas, E.P., and Raff, J.W. (2007). Maintaining the proper connection between the centrioles and the pericentriolar matrix requires Drosophila Centrosomin. J Cell Biology 178, 725–732.

Martinez-Campos, M., Basto, R., Baker, J., Kernan, M., and Raff, J.W. (2004). The Drosophila pericentrin-like protein is essential for cilia/flagella function, but appears to be dispensable for mitosis. J Cell Biology 165, 673–683.

Mattie, F.J., Stackpole, M.M., Stone, M.C., Clippard, J.R., Rudnick, D.A., Qiu, Y., Tao, J., Allender, D.L., Parmar, M., and Rolls, M.M. (2010). Directed microtubule growth, +TIPs, and kinesin-2 are required for uniform microtubule polarity in dendrites. Curr Biol 20, 2169–2177.

Meunier, S., and Vernos, I. (2016). Acentrosomal Microtubule Assembly in Mitosis: The Where, When, and How. Trends Cell Biol 26, 80–87.

Mitani, T., Punetha, J., Akalin, I., Pehlivan, D., Dawidziuk, M., Akdemir, Z.C., Yilmaz, S., Aslan, E., Hunter, J.V., Hijazi, H., et al. (2019). Bi-allelic Pathogenic Variants in TUBGCP2 Cause Microcephaly and Lissencephaly Spectrum Disorders. Am J Hum Genetics.

Munro, S. (2011). The Golgin Coiled-Coil Proteins of the Golgi Apparatus. Csh Perspect Biol 3, a005256.

Nakamura, N., Rabouille, C., Watson, R., Nilsson, T., Hui, N., Slusarewicz, P., Kreis, T.E., and Warren, G. (1995). Characterization of a cis-Golgi matrix protein, GM130. J Cell Biology 131, 1715–1726.

Nguyen, M.M., Stone, M.C., and Rolls, M.M. (2011). Microtubules are organized independently of the centrosome in Drosophila neurons. Neural Dev 6, 38.

Nguyen, M.M., McCracken, C.J., Milner, E.S., Goetschius, D.J., Weiner, A.T., Long, M.K., Michael, N.L., Munro, S., and Rolls, M.M. (2014). γ-Tubulin controls neuronal microtubule polarity independently of Golgi outposts. Mol Biol Cell 25, 2039–2050.

Ori-McKenney, K.M., Jan, L.Y., and Jan, Y.-N. (2012). Golgi outposts shape dendrite morphology by functioning as sites of acentrosomal microtubule nucleation in neurons. Neuron 76, 921–930.

Poirier, K., Lebrun, N., Broix, L., Tian, G., Saillour, Y., Boscheron, C., Parrini, E., Valence, S., Pierre, B.S., Oger, M., et al. (2013). Mutations in TUBG1, DYNC1H1, KIF5C and KIF2A cause malformations of cortical development and microcephaly. Nat Genet 45, 639–647.

Rao, A.N., Patil, A., Black, M.M., Craig, E.M., Myers, K.A., Yeung, H.T., and Baas, P.W. (2017). Cytoplasmic Dynein Transports Axonal Microtubules in a Polarity-Sorting Manner. Cell Reports 19, 2210–2219.

Rios, R.M. (2014). The centrosome-Golgi apparatus nexus. Philosophical Transactions Royal Soc B Biological Sci 369, 20130462–20130462.

Ríos, R.M., Sanchís, A., Tassin, A.M., Fedriani, C., and Bornens, M. (2004). GMAP- 210 Recruits γ-Tubulin Complexes to cis-Golgi Membranes and Is Required for Golgi Ribbon Formation. Cell 118, 323–335.

Rolls, M.M., and Jegla, T.J. (2015). Neuronal polarity: an evolutionary perspective. J Exp Biology 218, 572–580.

Roostalu, J., and Surrey, T. (2017). Microtubule nucleation: beyond the template. Nat Rev Mol Cell Bio 91, 321–710.

Roubin, R., Acquaviva, C., Chevrier, V., Sedjaï, F., Zyss, D., Birnbaum, D., and Rosnet, O. (2013). Myomegalin is necessary for the formation of centrosomal and Golgi-derived microtubules. Biol Open 2, 238–250.

Sanchez, A.D., and Feldman, J.L. (2016). Microtubule-organizing centers: from the centrosome to non-centrosomal sites. Curr Opin Cell Biol 44, 93–101.

Sánchez-Huertas, C., Freixo, F., Viais, R., Lacasa, C., Soriano, E., and Lüders, J. (2016). Non-centrosomal nucleation mediated by augmin organizes microtubules in post-mitotic neurons and controls axonal microtubule polarity. Nat Commun 7, 12187.

Schroeder, H.W., Hendricks, A.G., Ikeda, K., Shuman, H., Rodionov, V., Ikebe, M., Goldman, Y.E., and Holzbaur, E.L.F. (2012). Force-dependent detachment of kinesin-2 biases track switching at cytoskeletal filament intersections. Biophys J 103, 48–58.

Sears, J.C., and Broihier, H.T. (2016). FoxO regulates microtubule dynamics and polarity to promote dendrite branching in Drosophila sensory neurons. Dev Biol 418, 40–54.

Sepp, K.J., and Auld, V.J. (2003). Reciprocal Interactions between Neurons and Glia Are Required for Drosophila Peripheral Nervous System Development. J Neurosci 23, 8221–8230.

Stiess, M., Maghelli, N., Kapitein, L.C., Gomis-Rüth, S., Wilsch-Bräuninger, M., Hoogenraad, C.C., Tolić-Nørrelykke, I.M., and Bradke, F. (2010). Axon Extension Occurs Independently of Centrosomal Microtubule Nucleation. Science 327, 704–707.

Stone, M.C., Roegiers, F., and Rolls, M.M. (2008). Microtubules have opposite orientation in axons and dendrites of Drosophila neurons. Mol Biol Cell 19, 4122–4129.

Tas, R.P., Chazeau, A., Cloin, B.M.C., Lambers, M.L.A., Hoogenraad, C.C., and Kapitein, L.C. (2017). Differentiation between Oppositely Oriented Microtubules Controls Polarized Neuronal Transport. Neuron 96, 1264–1271.e5.

Teixidó-Travesa, N., Roig, J., and Lüders, J. (2012). The where, when and how of microtubule nucleation - one ring to rule them all. J Cell Sci 125, 4445 4456.

Torosantucci, L., Luca, M.D., Guarguaglini, G., Lavia, P., and Degrassi, F. (2008). Localized RanGTP accumulation promotes microtubule nucleation at kinetochores in somatic mammalian cells. Mol Biol Cell 19, 1873–1882.

Torres, I.L., Rosa-Ferreira, C., and Munro, S. (2014). The Arf family G protein Arl1 is required for secretory granule biogenesis in Drosophila. J Cell Sci 127, 2151–2160.

Tovey, C.A., and Conduit, P.T. (2018). Microtubule nucleation by γ-tubulin complexes and beyond. Essays Biochem 91, EBC20180028.

Tovey, C.A., Tubman, C.E., Hamrud, E., Zhu, Z., Dyas, A.E., Butterfield, A.N., Fyfe, A., Johnson, E., and Conduit, P.T. (2018). γ-TuRC Heterogeneity Revealed by Analysis of Mozart1. Curr Biol 28, 2314–2323.e6.

Triclin, S., Inoue, D., Gaillard, J., Htet, Z.M., Santis, M.D., Portran, D., Derivery, E., Aumeier, C., Schaedel, L., John, K., et al. (2018). Self-repair protects microtubules from their destruction by molecular motors. Biorxiv 499020.

Vinogradova, T., Miller, P.M., and Kaverina, I. (2009). Microtubule network asymmetry in motile cells: role of Golgi-derived array. Cell Cycle 8, 2168–2174.

Wang, Z., Wu, T., Shi, L., Zhang, L., Zheng, W., Qu, J.Y., Niu, R., and Qi, R.Z. (2010). Conserved Motif of CDK5RAP2 Mediates Its Localization to Centrosomes and the Golgi Complex. J Biol Chem 285, 22658–22665.

Weiner, A.T., Lanz, M.C., Goetschius, D.J., Hancock, W.O., and Rolls, M.M. (2016). Kinesin-2 and Apc function at dendrite branch points to resolve microtubule collisions. Cytoskeleton 73, 35–44.

Weiner, A.T., Seebold, D.Y., Torres-Gutierrez, P., Folker, C., Swope, R.D., Kothe, G.O., Stoltz, J.G., Zalenski, M.K., Kozlowski, C., Barbera, D.J., et al. (2020). Endosomal Wnt signaling proteins control microtubule nucleation in dendrites. Plos Biol 18, e3000647.

Wu, J., de Heus, C., Liu, Q., Bouchet, B.P., Noordstra, I., Jiang, K., Hua, S., Martin, M., Yang, C., Grigoriev, I., et al. (2016). Molecular Pathway of Microtubule Organization at the Golgi Apparatus. Dev Cell 39, 44 60.

Yadav, S., Younger, S.H., Zhang, L., Thompson-Peer, K.L., Li, T., Jan, L.Y., and Jan, Y.N. (2019). Glial ensheathment of the somatodendritic compartment regulates sensory neuron structure and activity. Proc National Acad Sci 116, 201814456.

Yalgin, C., Ebrahimi, S., Delandre, C., Yoong, L.F., Akimoto, S., Tran, H., Amikura, R., Spokony, R., Torben-Nielsen, B., White, K.P., et al. (2015). Centrosomin represses dendrite branching by orienting microtubule nucleation. Nat Neurosci 18, 1437–1445.

Yamada, M., and Hayashi, K. (2019). Microtubule nucleation in the cytoplasm of developing cortical neurons and its regulation by brain-derived neurotrophic factor. Cytoskeleton 76, 339–345.

Yan, J., Chao, D.L., Toba, S., Koyasako, K., Yasunaga, T., Hirotsune, S., and Shen, K. (2013). Kinesin-1 regulates dendrite microtubule polarity in Caenorhabditis elegans. Elife 2, e00133.

Yang, S.Z., and Wildonger, J. (2019). Golgi outposts locally regulate microtubule orientation in neurons but are not required for the overall polarity of the dendritic cytoskeleton. Biorxiv 866574.

Yau, K.W., Beuningen, S.F.B. van, Cunha-Ferreira, I., Cloin, B.M.C., Battum, E.Y. van, Will, L., Schätzle, P., Tas, R.P., Krugten, J. van, Katrukha, E.A., et al. (2014). Microtubule Minus-End Binding Protein CAMSAP2 Controls Axon Specification and Dendrite Development. Neuron 82, 1058–1073.

Zheng, Y., Wildonger, J., Ye, B., Zhang, Y., Kita, A., Younger, S.H., Zimmerman, S., Jan, L.Y., and Jan, Y.-N. (2008). Dynein is required for polarized dendritic transport and uniform microtubule orientation in axons. Nat Cell Biol 10, 1172–1180.

Zhou, W., Chang, J., Wang, X., Savelieff, M.G., Zhao, Y., Ke, S., and Ye, B. (2014). GM130 is required for compartmental organization of dendritic golgi outposts. Curr Biol 24, 1227–1233.

Zimyanin, V.L., Belaya, K., Pecreaux, J., Gilchrist, M.J., Clark, A., Davis, I., and Johnston, D.S. (2008). In vivo imaging of oskar mRNA transport reveals the mechanism of posterior localization. Cell 134, 843–853.

